# System-wide analyses of the fission yeast poly(A)+ RNA interactome reveal insights into organisation and function of RNA-protein complexes

**DOI:** 10.1101/748194

**Authors:** Cornelia Kilchert, Tea Kecman, Emily Priest, Svenja Hester, Krzysztof Kus, Alfredo Castello, Shabaz Mohammed, Lidia Vasiljeva

## Abstract

Production, function, and turnover of mRNA are orchestrated by multi-subunit machineries that play a central role in gene expression. Within these molecular machines, interactions with the target mRNA are mediated by RNA-binding proteins (RBPs), and the accuracy and dynamics of these RNA-protein interactions are essential for their function. Here, we show that fission yeast whole cell poly(A)+ RNA-protein crosslinking data provides system-wide information on the organisation and function of the RNA-protein complexes. We evaluate relative enrichment of cellular RBPs on poly(A)+ RNA to identify interactors with high RNA-binding activity and provide key information about the RNA-binding properties of large multi-protein complexes, such as the mRNA 3’ end processing machinery (cleavage and polyadenylation factor, CPF) and the RNA exosome. We demonstrate that different functional modules within CPF differ in their ability to interact with RNA. Importantly, we reveal that CPF forms additional contacts with RNA via the Fip1 subunit of the polyadenylation module and two subunits of the nuclease module. In addition, our data highlights the central role of the RNA helicase Mtl1 in RNA degradation by the exosome as mutations in Mtl1 lead to disengagement of the exosome from RNA. We examine how routes of substrate access to the complex are affected upon mutation of exosome subunits. Our results provide important insights into how different components of the exosome contribute to engagement of the complex with substrate RNA. Overall, our data uncover how multi-subunit cellular machineries interact with RNA, on a proteome-wide scale.

## Introduction

One of the major challenges in the field of RNA regulation is to understand how large multi-subunit molecular machineries interact with RNA. Crosslinking of recombinant *in vitro* reconstituted RNA-protein complexes has been a powerful tool to identify proteins that directly interact with RNA in the context of large RNP assemblies (Schmidt et al., 2012). However, this method requires protein complex purification, which in the case of large machineries is not a trivial task. Moreover, reconstituted complexes may not behave as in the intracellular environment, where they engage in active and dynamic interactions with RNA. These challenges prevent us from understanding how essential machineries that are involved in various aspects of RNA regulation such as RNA-processing and turnover function *in vivo*. In recent years, interactions of proteins with RNA have been systematically studied using *in vivo*, system-wide approaches. UV-crosslinking combined with protein purification has been used to identify RNA species bound by an RBP of interest (CLIP/CRAC) (Granneman et al., 2009; Hafner et al., 2010; Ule et al., 2003). Although these methods allow efficient identification of target RNAs and the specific sites where RBPs bind, the information is limited to individual proteins and cannot be used to compare in relative terms the RNA-binding activity of the subunits of a given complex. Recent development of UV-crosslinking-based methods in conjunction with oligo-d(T) enrichment followed by quantitative mass spectrometry (RNA Interactome Capture, RIC) has enabled to catalogue the “RBPome” of all polyadenylated RNAs in the cell in different model systems (Baltz et al., 2012; Beckmann et al., 2015; Castello et al., 2012; Conrad et al., 2016; Kwon et al., 2013; Marondedze et al., 2019; Matia-González et al., 2015; Mitchell et al., 2013; Sysoev et al., 2016). Here, we harness these technological advances to determine the RNA interactome of the fission yeast *Schizosaccharomyces pombe* (*S. pombe*). We show that by using whole cell extract (WCE)-normalisation of RIC, we can evaluate relative enrichment of cellular RBPs to assess the RNA-binding activity of individual proteins compared to other cellular proteins, and provide key information about the RNA-binding properties of large multiprotein complexes. We elucidate RNA interactions within the mRNA 3’ end processing machinery and the RNA exosome, which are involved in regulation of processing and stability of cellular RNA.

The 3’ ends of mRNAs, and most non-coding RNAs that are transcribed by RNA polymerase II (Pol II), are processed by the cleavage and polyadenylation machinery. This multi-subunit assembly consists of the cleavage and polyadenylation factor (CPF), and the accessory cleavage factors IA and IB (CFIA and CFIB). Additionally, we and others have recently identified an additional conserved factor (fission yeast Seb1 and human SCAF4) that associates with the 3’end processing machinery and is required for its function in fission yeast and humans (Gregersen et al., 2019; Kecman et al., 2018; Lemay et al., 2016; Wittmann et al., 2017). The mRNA 3’ end processing machinery recognizes polyadenylation sites (PAS) (AAUAAA in fission yeast (Mata, 2013; Schlackow et al., 2013) and humans (Proudfoot, 2011; Proudfoot and Brownlee, 1976)) and auxiliary regulatory sequences, and mediates cleavage and polyadenylation at the 3’ ends of transcripts (Thore and Fribourg, 2019; Zhao et al., 1999). Recent studies of the *S. cerevisiae* complex demonstrated that CPF is organised into three functionally different, stably associated modules: poly(A) polymerase, nuclease, and phosphatase (Casañal et al., 2017). Reconstitution and structural analysis of the mammalian polymerase module (CPSF160-WDR33-CPSF30-Fip1) have revealed that the zinc finger domains 2 and 3 of CPSF30, as well as the WD40 domain and the N-terminal region of WDR33, interact with the PAS element (Clerici et al., 2017, 2018; Schönemann et al., 2014; Sun et al., 2018). Although RNA is not present in the cryo-EM structure of the *S. cerevisiae* polymerase module, its overall organisation is very similar to the mammalian module (Casañal et al., 2017). The residues involved in PAS recognition by CPSF30 and the WD40 domain of WDR33 (but not its N-terminus) are conserved in their *S. cerevisiae* homologues Yth1 and Pfs2, suggesting that these subunits might bind PAS in yeast as well. At the same time, other subunits of the polymerase module, CPSF160/Cft1 and hFip1/Fip1, are not in contact with the PAS in the structure and were proposed to act as scaffolding proteins. Despite this progress, it is not clear how interactions between the pre-mRNA and the full complex apart from PAS recognition help to ensure accuracy of PAS site selection during 3’ end processing. In agreement with recent structural data, we show that each functional module of CPF differs in its RNA-binding behaviour. Our data support the proposed role of the polyadenylation module in PAS recognition. Importantly, we show that CPF forms additional contacts with RNA, which are mediated by Fip1 and components of the nuclease module. In contrast, the phosphatase module is not enriched on RNA.

We also applied RIC to study RNA interactions of another important cellular machinery, the RNA exosome and its co-factors (Kilchert et al., 2016; Morton et al., 2018). The exosome complex is composed of a nine subunit barrel-like structure, which is associated with the 3’-5’ exonucleases Rrp6 and Dis3 located at the top and bottom, respectively (Makino et al., 2013, 2015; Schuch et al., 2014; Zinder et al., 2016). RNA molecules are threaded through the central channel in 3’ to 5’ direction before they are degraded by Dis3, which also has endonucleolytic activity (Lebreton et al., 2008; Schaeffer et al., 2009; Schneider et al., 2009). Although the exosome complex has important functions in controlling levels of various types of cellular RNAs, the mechanisms underlying its regulation are poorly understood. We show that exosome co-factors are RBPs and propose that interaction with RNA is important for their function in exosome regulation. We find that mutation of the essential Ski2-like RNA helicase Mtl1 leads to disengagement of the exosome from RNA, which supports a model where Mtl1 is central to mediating RNA degradation within the exosome. Furthermore, using a comparative RIC approach, we monitor changes in the preferred route of substrate access to the RNA exosome complex in different exosome mutants and provide important insights into how various exosome subunits contribute to the interaction of the complex with substrate RNA.

## Results and discussion

### *S. pombe* poly(A)+ RNA interactome capture

Fission yeast recapitulates many aspects of mammalian RNA regulation. Yet, no systematic analysis had been performed to identify proteins that bind RNA and could potentially play a role in RNA regulation in this important model organism. For this reason, we determined the *S. pombe* poly(A)+ RNA interactome. To facilitate RNA-protein UV-crosslinking, wild-type cells were labelled with 4-thiouracil (4sU). RNPs were enriched by oligo-d(T) selection and RNA-associated proteins identified by mass spectrometry (MS). We discovered 805 proteins significantly enriched over the non-irradiated control (n=6; p < 0.01) (Figure 1A-C, S1A). As expected, gene ontology (GO) analysis revealed RNA-related GO terms to be significantly enriched in our *S. pombe* RBPome, including “RNA metabolic process” (p = 8.63·10^-27^) and “RNA binding” (p = 4.58·10^-63^). We rediscovered 99 of 136 proteins annotated to harbour a classical RNA-binding domain (RBD) in *S. pombe* (e.g. RRM, DEAD, KH; see Supplemental Table S1 for a complete list; p < 0.01), highlighting the depth our RBPome and the presence of the expected molecular signatures (Figure 1C).

**Figure 1.**
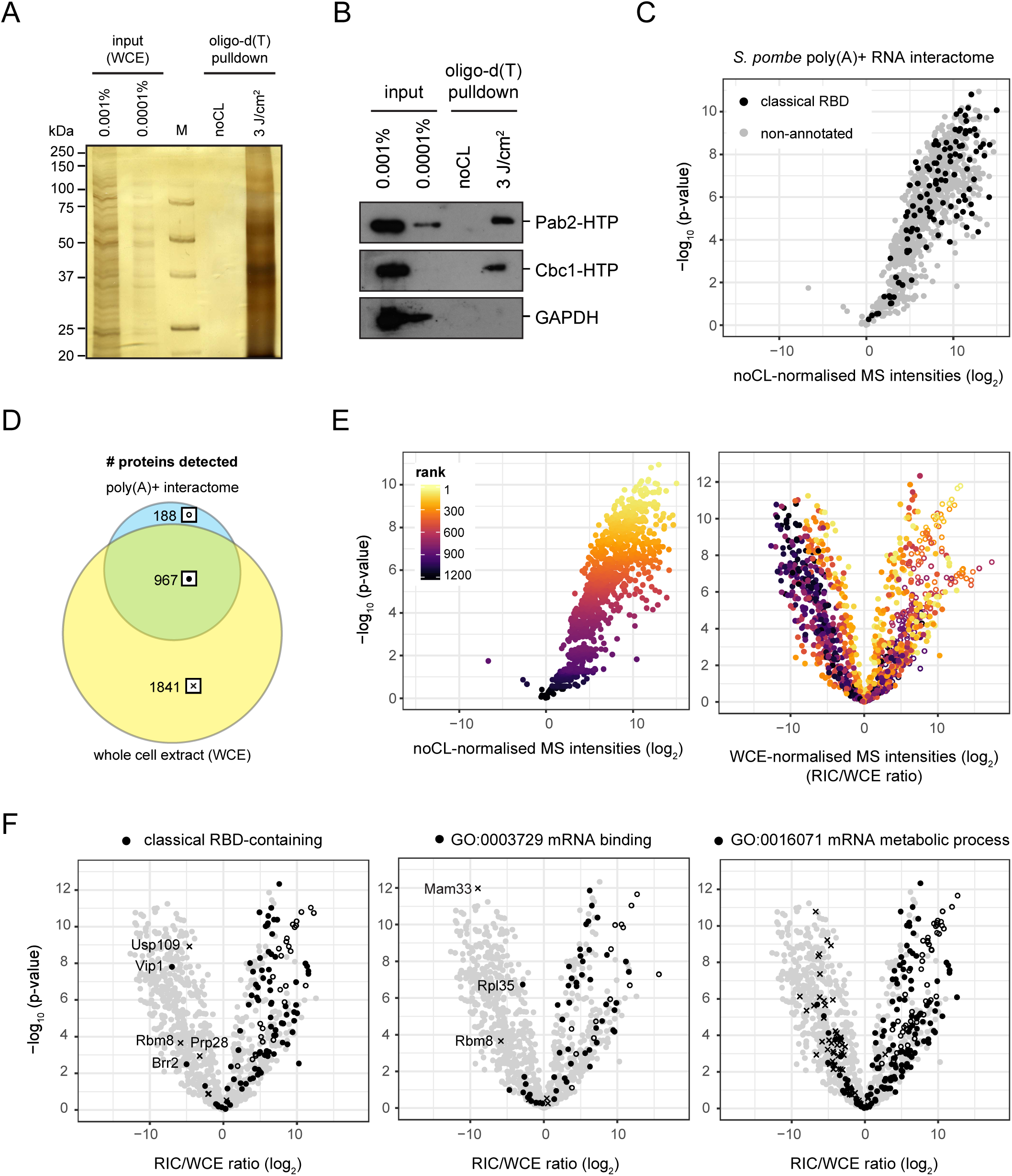
poly(A)+ RNA interactome capture in *S. pombe.* Quality control of the RNA interactome capture using silver staining **(A)** or Western blotting **(B)** of input samples and RNA interactors eluted from oligo-dT beads. Due to stringent washes, RNA interactors are only recovered if cells are irradiated with UV prior to harvesting (3 J/cm^2^), but not from the non-crosslinked controls (noCL). Known RBPs such as the nuclear poly(A)-binding protein Pab2 or the cap-binding protein Cbc1 are robustly detected in the crosslinked samples, whereas the abundant metabolic enzyme GAPDH (glyceraldehyde-3-phosphate dehydrogenase) is not enriched. **(C)** Mass spectrometry (MS) analysis of the *S. pombe* poly(A)+ RNA interactome. In the volcano plot, p-values (-log, moderated Student’s t-test) are plotted against the fold change of mean MS intensities (log_2_) of proteins recovered from the oligo-d(T) pull-downs of UV-crosslinked samples (3 J/cm^2^) normalised to the non-crosslinked controls (noCL) (n=6). Background values were imputed for proteins without signal in the noCL control. Proteins annotated with a classical RNA-binding domain (RBD) are designated in black. The complete list of Pfam identifiers used for this analysis can be found in Supplemental Table 1. 805 proteins were significantly enriched in the UV-crosslinked samples (p-value ≤ 0.01). **(D)** Overlap of proteins detected in oligo-d(T) pull-downs of UV-crosslinked wild-type cells (3 J/cm^2^) and the corresponding whole cell extracts (WCE, n=6). WCE coverage of proteins detected in the pull-down was 83.7%. In downstream analyses, the indicated symbols are used to designate proteins from a given population: full circles for proteins detected in both oligo-d(T) pull-down and WCE; empty circles for proteins detected in the oligo-d(T) pull-downs but not the WCE; crosses for proteins detected in the WCE but not the oligo-d(T) pull-downs. **(E)** Impact of normalisation method on relative RBPs enrichment in the poly(A)+ RNA interactome. In these volcano plots, p-values (-log, moderated Student’s t-test) are plotted against the fold change of mean MS intensities (log_2_) of proteins recovered from the oligo-d(T) pull-downs of UV-crosslinked samples (3 J/cm^2^) over either the non-crosslinked controls (noCL, left panel) or the input WCE (RIC/WCE ratios, right panel) (n=6). In the right panel, background values were imputed for proteins without WCE signal (empty circles). For WCE normalisation (right panel), raw MS intensities were normalised to median = 0 for all samples prior to calculating fold change values. In both panels, individual proteins are coloured according to the statistical significance (ranked p-values) of protein enrichment in the noCL-normalised interactome. **(F)** Distribution of proteins annotated with a classical RBD (left panel), GO function “mRNA binding” [GO:0003729] (middle panel) or GO process “mRNA metabolic process” [GO:0016071] (right panel) in the WCE-normalised RNA interactome, indicated in black. Full circles denote proteins that were detected in both oligo-d(T) pull-down and WCE, empty circles proteins that were present in the oligo-d(T) pull-downs but not the WCE, crosses proteins that were detected in the WCE, but never in the oligo-d(T) pull-downs (see also 1C). Background values were imputed for proteins without signal in any given sample.

### Enrichment over WCE reflects degree of poly(A)+ RNA association

We hypothesized that referencing our data to cellular protein abundances would allow to better assess the relative RNA-binding activity of the identified RBPs. Recent studies that compare RNA interactomes under different conditions have employed this approach to determine whether observed changes in RNA binding were a consequence of changes in RBP abundance (Garcia-Moreno et al., 2019; Sysoev et al., 2016). For this, we analysed the abundance of individual proteins in the whole cell extracts (WCE). WCE MS intensities were then used to normalise the RIC data for protein abundance (Figure 1D, E and S1B). 83.7% of all proteins identified by RIC were also detected in the WCE, that thus had an excellent coverage. Background values were imputed for proteins that were not detected in the WCE sample (Figure S1C and S1D). WCE-normalisation revealed that many proteins with high MS intensities in the RIC experiment were poorly captured considering their abundance. By contrast, many proteins with low to intermediate signal in RIC were captured very efficiently as they were of low abundance in the WCE (Figure 1E). We expect RIC/WCE ratios to reflect a combination of two RBP characteristics: i) the fraction of the RBP that associates with poly(A)+ RNA *in vivo*, i.e. its RNA-binding activity, which is determined by the affinity of the RBP for its target RNA and target RNA availability; ii) the UV-crosslinkability of RBP and the bound RNA, which is determined by the geometry of the interaction (the spatial configuration of amino acids and nucleotide bases), and also the type of residues present at the interface.

To estimate the relative impact of UV-crosslinkability on RIC/WCE ratios, we considered proteins that are expected to have high poly(A)+ RNA-binding activity. Through an optimal interaction surface, classical RBDs are known to bind RNA with high affinity (Lunde et al., 2007). Strikingly, proteins with a classical RBD were almost uniformly enriched relative to other RNA interactors (Figure 1F, left panel). For abundant classical RBDs (≥ 4 separate annotated proteins in *S. pombe*), variability of RIC/WCE ratios was higher between RPBs with the same domain than between different RBDs, and we could not detect any significant influence of classical RBD type on enrichment in RIC (Figure S1E and F, p > 0.01). When comparing RBPs with one or multiple domains, the degree of enrichment in the RBPome did not markedly increase with the number of annotated classical RBDs (Figure S1G).

Significant enrichment was also observed for most proteins known to have high specificity for poly(A)+ RNA based on GO term annotation, for example “mRNA binding” or “mRNA metabolic process” (Figure 1F, middle and right panel). At the same time, 4sU-dependent UV-crosslinking was specific to RNA: For example, annotated RNA helicases showed high average enrichment in RIC. In contrast, annotated DNA helicases rarely crosslinked, and only two proteins, Spcc737.07c and Snf22, were significantly enriched (Figure S2A), possibly indicating that these are helicases with dual specificity.

Poor UV-crosslinkability can lead to an underestimation of the degree of RNA association for RBPs with high *in vivo* poly(A)+ RNA-binding activity. To estimate the number of false-negative data points that arise from this, we looked for known mRNA binders that failed to be enriched in the RIC experiment. 99 classical RBPs and 55 “mRNA binding” proteins were robustly detected in RIC. In contrast, 14 and 6 RBPs, respectively, were detected but not significantly enriched compared to the non-crosslinked control (p > 0.01; Figure 1C and data not shown), while 5 and 7 RBPs, respectively, never crosslinked at all although they were present in the WCE (Figure 1F, left and middle panels, indicated by crosses).

Strikingly, 10 out of 19 classical RBPs that were RIC-negative are known to be associated with non-poly(A) RNA species (e.g. Usp109 and Brr2 / Prp28 with U1 and U5 snoRNPs, respectively; Figure 1F, left panel) or involved in ribosome biogenesis. Four others are meiosis-specific RBPs, or mitochondrial proteins. Importantly, mitochondrially encoded transcripts fail to be enriched after oligo-d(T) selection in *S. pombe* as they are not polyadenylated (Marguerat et al., 2012).

Similarly, 30% of all RIC-negative “mRNA binding” proteins are mitochondrial RBPs, for example Mam33, the homologue of a nuclear-encoded mitochondrial translation regulator in budding yeast (Roloff and Henry, 2015) (Figure 1F, middle panel). Rbm8/Y14, on the other hand, is a component of the exon-exon junction complex that binds spliced mRNA, but - at least in the human complex - the RNA-binding surface of the RBD is completely masked by Magoh (Lau et al., 2003). For other proteins, such as Vip1, the mode of binding and the nature of the bound RNAs are unknown.

Among proteins annotated with “mRNA metabolic process”, we found several RNP complexes that were inefficiently captured with oligo-d(T), including CCR4-NOT deadenylase and the transcription elongation factor complex (Figure S2B and 3B), which bind deadenylated mRNA or nascent RNA still lacking a poly(A) tail, respectively. In contrast, RBPs known to interact with fully processed mRNA are almost uniformly enriched. In summary, we observe robust enrichment of RBPs that are expected to associate with poly(A)+ RNA based on annotation, although these RBPs must be assumed to UV-crosslink with a whole range of different efficiencies. We take this as an indication that differences in UV-crosslinkability – although they modulate RIC/WCE ratios – do not have a dominant influence on RIC/WCE values, and that the false-negative rate for RIC is reasonably low.

Because of the unknown UV-crosslinkability component, RIC/WCE ratios cannot be regarded as actual measurements of *in vivo* poly(A)+ RNA-binding activity. However, we conclude that RIC/WCE ratios can serve as an estimator of the degree of *in vivo* poly(A)+ RNA association with a certain confidence. In the following, we demonstrate the uses of this assumption.

### Classification of non-classical RBDs based on RIC/WCE ratios

First, we sought to gain additional information on RBPs that lack classical RBDs. Recent studies have greatly expanded the scope of RBPs that interact with RNA (Baltz et al., 2012; Beckmann et al., 2015; Castello et al., 2012, 2016; Kwon et al., 2013; Matia-González et al., 2015; Mitchell et al., 2013). These proteins harbour domains with not well-established modes of RNA binding. Moreover, a recent method, RBDmap, has adapted the RIC workflow to identify the regions of RBPs engaged in RNA binding (Castello et al., 2016). Together, annotated non-canonical RBDs and RNA-binding regions discovered by RBDmap comprise 192 Pfam identifiers (Castello et al., 2012, 2016; Moore et al., 2018) (Supplemental Table S1), where 148 of these have annotations in *S. pombe*. If analysed conjointly, the behaviour of non-classical RBD-containing proteins differs strikingly from classical RBPs: Many non-classical RBPs have low RIC/WCE ratios, indicating that they are present in the RNA interactome, but strongly underrepresented (Figure 2A, right panel). We conclude that – in stark contrast to classical RBPs – many RBPs with non-classical RBDs that were defined based on UV-crosslinking experiments have low RNA-binding activity. We expect these to associate with poly(A)+ RNA at substoichiometric levels under standard growth conditions (Figure 2B). However, non-classical RBPs display a broad range of RIC/WCE ratios, from very high to very low RNA-binding activity. This can be visualised when RIC/WCE ratios are plotted according to non-classical RBD annotation (Figure S3). The number of annotated proteins for most non-classical RBDs in *S. pombe* is too low to allow reliable assignment of an average RNA-binding activity. Thus, we focused on only those RBDs present in at least four different proteins detected in our RIC experiment (Figure S3). We can distinguish: i) domains harboured by proteins with high RIC/WCE ratios, which we refer to as “classical-like”, and include LSM, S1 and C2H2-type zinc finger domains. These are likely to represent professional RBDs that are nearly constantly associated with RNA. ii) Domains harboured by proteins with low RIC/WCE ratio, that we refer to as “substoichiometric”, and include TCP-1/cpn60 chaperonin family proteins, Hsp70 proteins, thioredoxins, aldehyde dehydrogenase family proteins, and cyclophilin-type peptidyl-prolyl cis-trans isomerases. iii) Domains harboured by proteins with a broad range of RIC/WCE ratios, which we refer to as “adaptive” RBDs, and include WD40 repeat proteins, Elongation factor Tu domain 2-containing proteins, Elongation factor Tu GTPase domain, 50S ribosome-binding GTPase and Helicase C domain-containing proteins. In the case of WD40 repeats, a common protein-protein interaction fold, the inclusion of basic amino acids in the binding surface was found to correlate with its ability to bind RNA (Castello et al., 2012, 2016).

**Figure 2.**
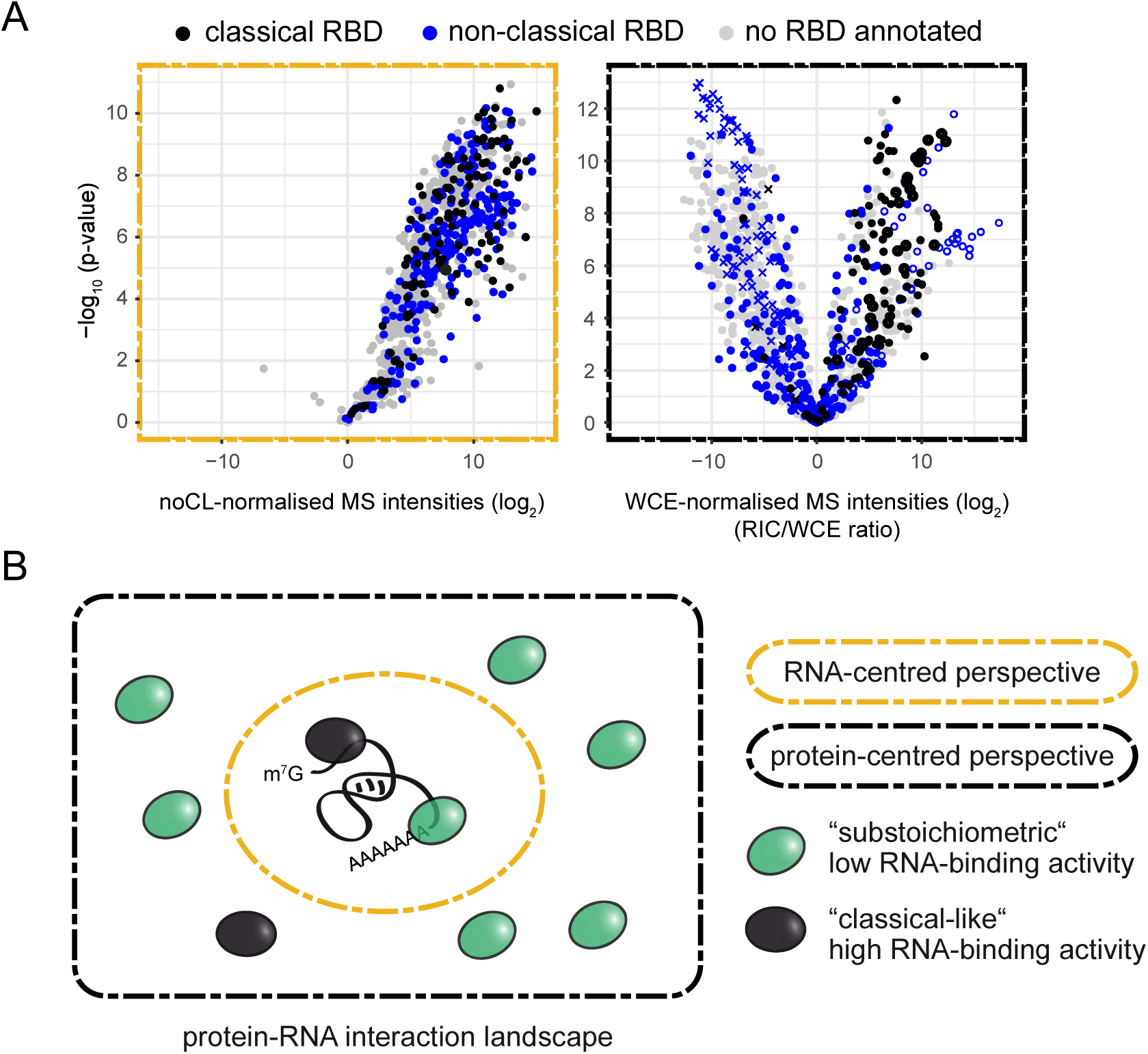
Properties of non-classical RBDs. **(A)** Distribution of proteins annotated with a classical or non-classical RBD in the noCL-normalised (left panel) and WCE-normalised poly(A)+ RNA interactome (right panel). Non-classical RBDs comprise 192 Pfam identifiers derived from the initial human interactome as well as any domain in RBDmap that had at least three peptides supporting it (Castello et al., 2012, 2016; Moore et al., 2018). The complete list of Pfam identifiers used for this analysis can be found in Supplemental Table S1. Full circles denote proteins that were detected in both oligo-d(T) pull-down and WCE, empty circles proteins that were present in the oligo-d(T) pull-downs but not the WCE, crosses proteins that were detected in the WCE, but never in the oligo-d(T) pull-downs (see also 1C). Background values were imputed for proteins without signal in any given sample. **(B)** The mode of normalisation determines the information extracted from RIC data: Normalisation to a non-crosslinked control will yield a quantitative snapshot of the partial proteome that an average RNA molecule interacts with (RNA-centred perspective, indicated in orange). In contrast, normalisation to protein abundances will yield the relative enrichment of RBPs in the interactome, thus providing an estimate of the fraction of an RBP that is associated with poly(A)+ RNA, i.e. of its poly(A)+ RNA-binding activity (protein-centred perspective, indicated in black). As the comparison between noCL-normalised and WCE-normalised RIC data reveals, in the sum, the number of molecular contacts between RNA and highly abundant substoichiometric RNA interactors rivals or even outweighs interactions with RBPs with high RNA-binding activity that are expressed at low levels.

RBPs with low RIC/WCE ratios are the most intriguing class. Due to the limitations of the poly(A)+ RNA selection procedure, some of these proteins may preferentially bind non-poly(A) RNA species – likely examples are Nhp2 and Snu13, which are components of box H/ACA and U3 snoRNPs, respectively, and annotated with PF01248 [ribosomal_7Ae]. Others may be activated under particular physiological conditions. RNA-binding of proline cis/trans isomerases and chaperones, for example, is triggered in response to virus infection (Garcia-Moreno et al., 2019). Others, however, may either bind to RNA with low affinity or display “moonlighting” RNA-binding activities. For example, mouse glyceraldehyde-3-phosphate dehydrogenase (GAPDH), a glycolytic enzyme, was previously reported to bind and translationally inhibit IFN-γ-mRNA in a manner that is inversely correlated with the glycolytic activity of the cell, and only a fraction of GAPDH molecules associates with RNA during normal growth (Chang et al., 2013). Another example is iron-responsive protein 1 (IRP1), also known as aconitase 1 (ACO1). Under low iron levels, a proportion of ACO1/IRP1 loses its iron-sulphur cluster and as apo-protein becomes an RBP (Muckenthaler et al., 2017). The list of enzymes that were confirmed to moonlight as RNA-binders and/or be regulated through an RNA-binding event has grown over the years (Castello et al., 2015; Cieśla, 2006). Befittingly, many of these are represented among the substoichiometric RNA interactors that we define based on RIC/WCE ratios. We suggest that RNA binding is a secondary function for most of these low RIC/WCE ratio RNA binders. In support of this, only 3.6% of proteins that contain a non-classical RBD and bind RNA in the substoichiometric range are annotated with the GO term “mRNA metabolic process”, which is close to the proteome-wide average of 6.2%. This is opposed to 13.1% of proteins annotated with an adaptive domain, and 30% of proteins that display classical-like strong RNA-binding behaviour. We are confident that the classification of RBPs based on their observed RIC/WCE ratio will be instrumental to predict the type of biological role they play.

### Characterisation of RBPs with high RNA-binding activity that lack annotated RBDs

In addition to known RBPs, we identified numerous RBPs with high RNA-binding activity that were not annotated with any classical or non-classical RBD from our list (Supplemental Table S2). This group of proteins is of high interest as it could include novel RBPs with unexpected functions in RNA regulation to be mined in the future. Interestingly, when comparing proteins without annotated RBDs that were enriched in the interactome (RIC/WCE ratio > 4, p < 0.01) to those that were underrepresented (RIC/WCE ratio < 0.25, p < 0.01), enriched proteins were more likely to be nuclear, to be involved in gene expression and DNA metabolic processes, and to be annotated as RNA-, DNA-, or metal-ion-binding, or to possess transcription regulator activity (Figure 3A). We were struck by the high representation of DNA- or transcription-related GO terms among highly active RNA binders. Recently, an increasing number of studies have reported cases of chromatin-associated proteins that are bound and/or regulated by RNA binding (Hendrickson et al., 2016). A prominent example is the PRC2 complex, a key chromatin modifier which binds RNA promiscuously via non-canonical RNA-binding elements that are dispersed across the surface of the complex (Cifuentes-Rojas et al., 2014; Davidovich et al., 2013; Kaneko et al., 2014; Long et al., 2017). In our *S. pombe* data set, the following proteins with GO function “histone modification” were significantly enriched on poly(A)+ RNA: the histone deacetylase complex subunit Sap18; the argonaut protein Ago1; and the rRNA methylase and potential histone H2A methylase Fib1 (fibrillarin). Interestingly, SPAC25G10.01, a homologue of TRRAP, which is a component of the histone acetylation complex in humans, is also highly enriched. Mutations in TRRAP have been found to be a cause of autism and syndromic intellectual disability in humans (Cogné et al., 2019). It is tempting to speculate that RNA binding might play a regulatory role in modulating the activity of the histone acetyltransferase complex, an exciting possibility that could be tested in the future.

**Figure 3.**
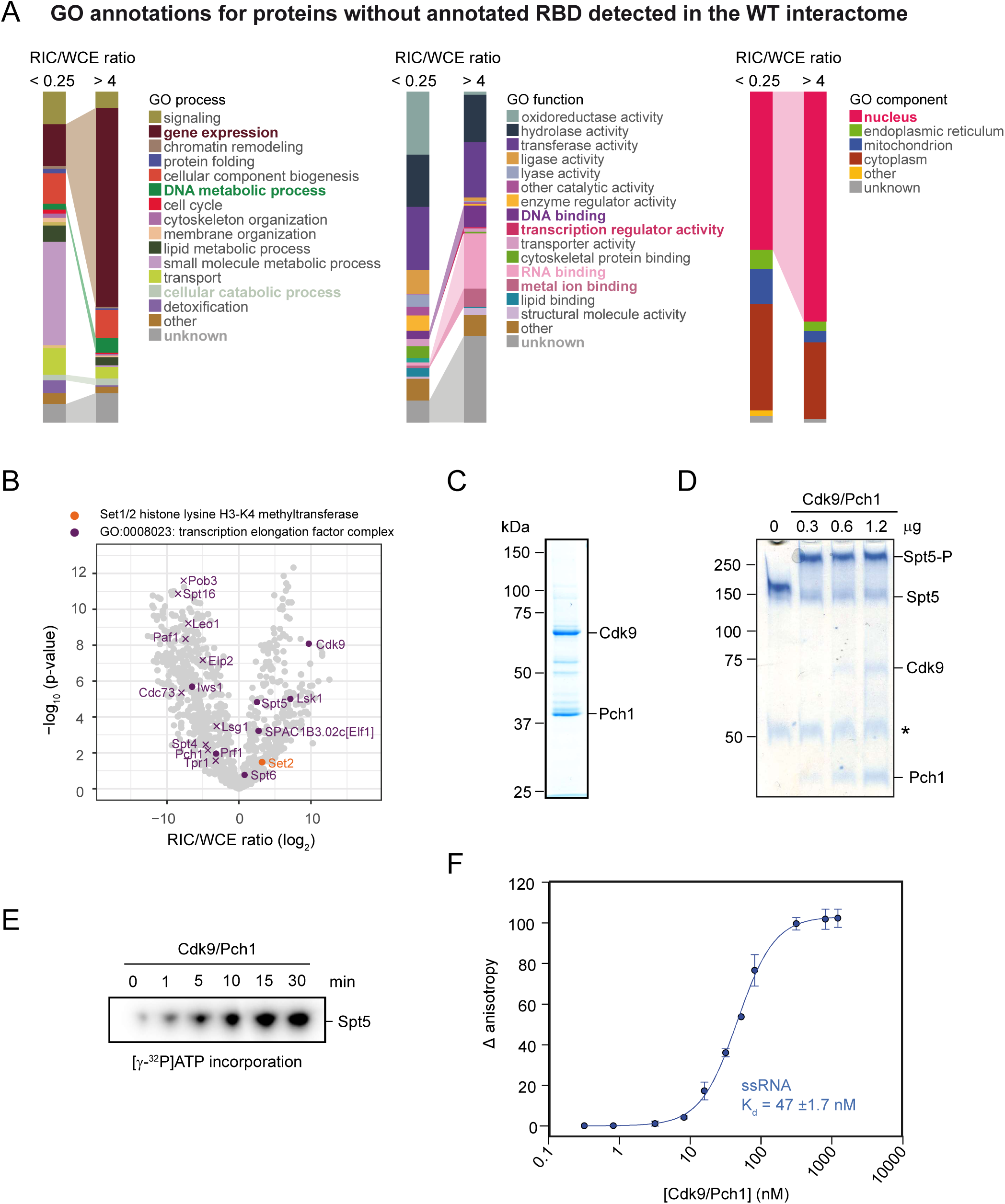
Characterisation of RBPs with high RNA-binding activity that lack annotated RBDs: The serine/threonine kinase Cdk9 binds single-stranded RNA with high affinity. **(A)** Comparison of GO term distribution for non-RBD-containing proteins that are either underrepresented (RIC/WCE ratio < 0.25; n = 304) or enriched (RIC/WCE ratio > 4; n = 203) in the poly(A)+ RNA interactome (p < 0.01), visualised with the QuiLT tool on PomBase (Lock et al., 2018). Categories overrepresented among proteins enriched in the WT interactome compared to other RNA interactors are highlighted in bold. **(B)** Distribution of proteins annotated with GO component “transcription elongation factor complex” [GO:0008023] in the WCE-normalised RNA interactome, indicated in purple. The histone lysine H3-K4 methyltransferase Set2 is indicated in orange. Set1 was not detected in any of the samples. Full circles denote proteins that were detected in both oligo-d(T) pull-down and WCE, empty circles proteins that were present in the oligo-d(T) pull-downs but not the WCE, crosses proteins that were detected in the WCE, but never in the oligo-d(T) pull-downs (see also 1C). Background values were imputed for proteins without signal in any given sample. **(C)** Coomassie stain of Cdk9/Pch1 purified from insect cells. **(D)** Phosphotag gel of in vitro kinase assays with increasing amounts of Cdk9/Pch1 on recombinantly purified Spt4/5. Phospho-tag changes the migration of phosphorylated proteins. The asterisk marks an unspecific band that copurifies with Spt5. **(E)** Autoradiogram of in vitro kinase assays of Cdk9/Pch1 with [γ-^32^P]ATP on recombinantly purified Spt4/5. **(F)** Fluorescence anisotropy assay measuring binding of increasing amounts of Cdk9/Pch1 to 8 nM FAM-labelled RNA. The observed Kd-value was 47 ±1.7 nM.

In addition, we found a number of “DNA-binding transcription factors” to be highly enriched on poly(A)+ RNA, including the shuttle craft/ NFX1 homologue Spcc18.03, and a calcineurin-responsive transcription factor, Prz1 (Figure S4A). The vast majority of proteins annotated as GO component “transcription elongation factor complex” were not identified by RIC. Surprisingly, several transcription elongation factors (TEFs), i.e. Spt5, Elf1, and the serine/threonine kinases Cdk9 and Lsk1, displayed high RIC/WCE ratios, while Spt6, the histone lysine H3-K36 methyltransferase Set2, and the Paf1 complex member Prf1 showed intermediate enrichment on poly(A)+ RNA (Figure 3B). In a recent CLIP study conducted on a panel of TEFs in *S. cerevisiae*, Spt5, Spt6, Ctk1/Lsk1, Bur1/Cdk9, Set2, and Rtf1/Prf1 were found to bind nascent pre-mRNAs (Battaglia et al., 2017). The strong enrichment of Spt5, Cdk9, and Lsk1 on poly(A)+ RNA was therefore unexpected. It suggested that these TEFs stay associated with mRNAs post-transcriptionally, and therefore might be interacting with RNA independently of the Pol II transcription machinery. In short, the identification of these three proteins by RIC implied that they directly bind RNA with high affinity.

Cdk9 (P-TEFb) is a cyclin-dependent protein kinase that controls early elongation of RNA Pol II (Price, 2000). When bound to the cyclin Pch1, Cdk9 phosphorylates various regulators of transcription elongation, including Spt5 (Pei et al., 2003). In *S. cerevisiae*, the cyclin-dependent Pol II kinase complex Ctk1/Ctk2/Ctk3 (Lsk1/Lsc1 in *S. pombe*) has been reported to bind RNA in the nanomolar range (Battaglia et al., 2017). To validate RNA binding of Cdk9 *in vitro*, we co-expressed full-length *S. pombe* Cdk9 together with its cognate cyclin Pch1 in insect cells (Fig. 3C). We then tested the purified Cdk9/Pch1 complex for kinase activity using a purified substrate, Spt5. Spt5 was efficiently phosphorylated by Cdk9/Pch1 *in vitro*, showing that the purified kinase complex was active (Fig. 3D and E). We then tested the ability of Cdk9/Pch1 to interact with RNA by fluorescence anisotropy (Fig. 3F).

Cdk9/Pch1 bound RNA with high affinity (Kd=47.0±1.7 nM). These *in vitro* results confirmed that Cdk9 strongly interacts with RNA as reported by RIC, and that our approach can identify novel RBPs that are characterised by high RNA-binding activity. Amongst annotated serine/threonine kinases, Cdk9 and Lsk1 were particularly enriched on poly(A)+ RNA. Interestingly, comparable poly(A)+ RNA-binding activities were observed for the atypical protein kinases Rio1/2 (Figure S4B). Rio kinases have a conserved role in rRNA maturation and were recently reported to be regulated by binding to rRNA (Knüppel et al., 2018). However, a broader role for Rio kinases in regulating nutrient-activated expression of genes has been proposed (Iacovella et al., 2018). Our data support a model where Rio kinase activity could be regulated by more diverse RNA species.

### Positioning the mRNA tunnel within ribosomes using RIC/WCE ratios

Because WCE-normalised RIC provided information about relative RNA-binding activities, we rationalised that RIC could be used to position RNA within large multi-protein complexes. To test this hypothesis, we analysed the enrichment of ribosomal proteins (RPs) on poly(A)+ RNA. In the ribosome, most structural RPs interact with non-poly(A) rRNA. By contrast, RPs that line the mRNA tunnel should be able to establish interactions with poly(A)+ mRNA. As expected, most RPs annotated as components of the “cytosolic ribosome” were captured with low efficiency. 15 RPs were detected in the WCEs but were absent from the RIC data set, suggesting that they do not interact, even stochastically, with poly(A)+ RNA. However, RPs that exhibited high recovery by RIC, such as S2 and S3, are immediately adjacent to the mRNA channel (Figure 4A and B, indicated in blue). Hence, the WCE-normalised RIC data correlates well with expectations for protein-poly(A)+ RNA association based on the known structure of the ribosome (Schmidt et al., 2016). We conclude that normalisation against the WCE adds critical information to the RIC data, and allows to make inferences about how large RNP complexes assemble on poly(A)+ RNA.

**Figure 4.**
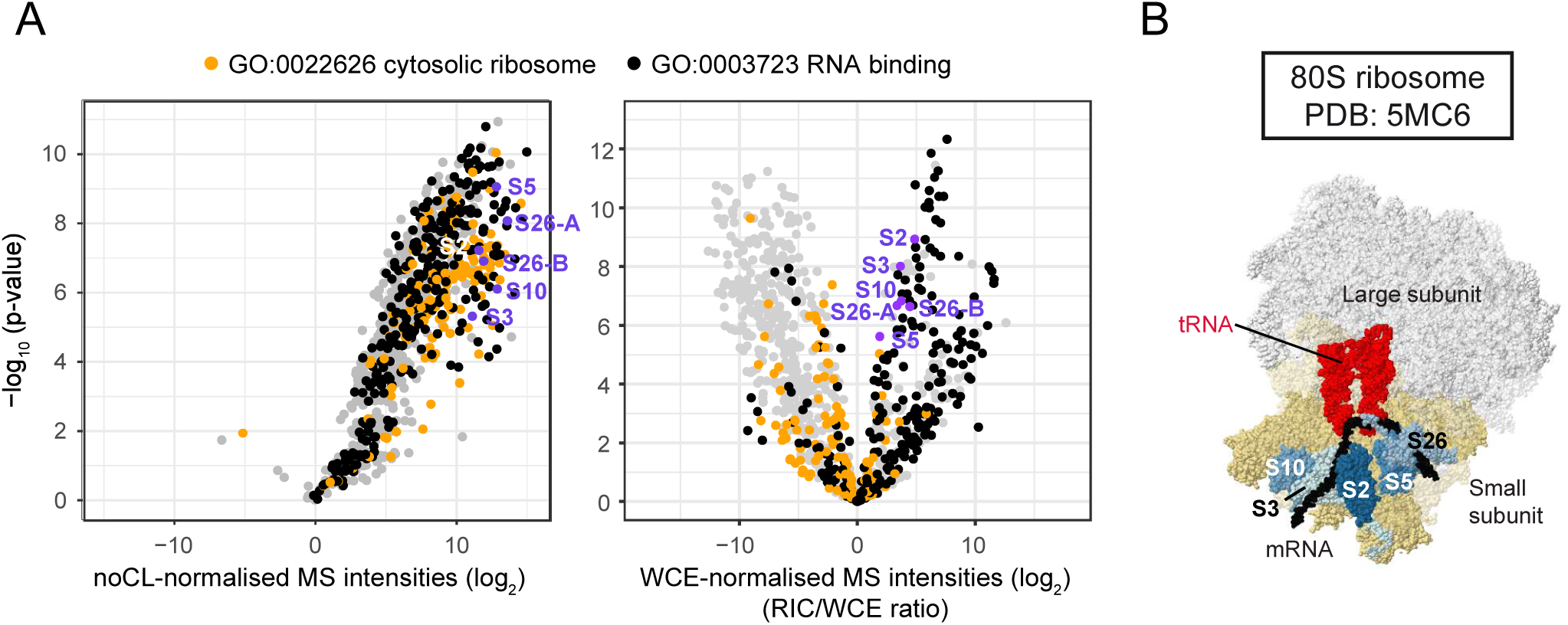
Positioning the mRNA tunnel within ribosomes using RIC/WCE-ratios. **(A)** Distribution of ribosomal proteins in the normalised poly(A)+ interactome. Volcano plot of the noCL-normalised (left panel) or WCE-normalised (right panel) RNA interactome as in 1D. Proteins annotated as cellular component “cytosolic ribosome” [GO:0022626] are designated in orange, with the molecular function “RNA binding” [GO:0003723] in black. For proteins annotated with both GO terms, GO:0022626 was given precedence. The ribosomal proteins that were most significantly enriched on poly(A)+ RNA were mapped to the *S. cerevisiae* ribosome structure **(B)** and are indicated in blue. Proteins without WCE signal were disregarded (see also Figure S4C). **(B)** Structure of the *S. cerevisiae* 80S ribosome [PDB:5MC6] (Schmidt et al., 2016). Ribosomal proteins with *S. pombe* orthologues that were highly significantly enriched on poly(A)+ RNA are located close to the mRNA channel in the structure, and are designated in blue. For reference, the mRNA is coloured black, tRNAs bound in the ribosomal A- and P-sites red, and the small and large ribosomal subunits beige and white, respectively.

### Using RIC to study the topology of multi-protein complexes

Due to its complex and dynamic nature, our understanding of how the mRNA 3’end processing machinery interacts with pre-mRNA is limited. To gain more information, we analysed the relative enrichment of subunits of the fission yeast CPF on poly(A)+ RNA (Figure 5A and B). Interestingly, components of the polymerase module, as well as the CFIB factor Msi2, crosslinked very well to poly(A)+ RNA, suggesting that these modules stay associated at least for some time with the polyadenylated mRNA following endonucleolytic cleavage. Consistent with the described role of Pfs2 and Yth1 in PAS recognition in humans, both factors were highly enriched on RNA in fission yeast, suggesting that their function within CPF is likely to be evolutionary conserved. Surprisingly, Fip1 crosslinked equally well, comparable to Pfs2 and Yth1, suggesting that it could form additional contacts with RNA contributing to either stability or specificity of the CPF-RNA interaction. In contrast, other subunits of the polymerase module, Ctf1 and the poly(A) polymerase Pla1, were not strongly enriched on poly(A)+ RNA, supporting the proposed role of Cft1 as a scaffold. Two components of the nuclease module of CPF, Mpe1 and Cft2 (but not Ipa1 or Ysh1 endonuclease), also interacted with RNA, possibly to assist in positioning Ysh1 nuclease.

**Figure 5.**
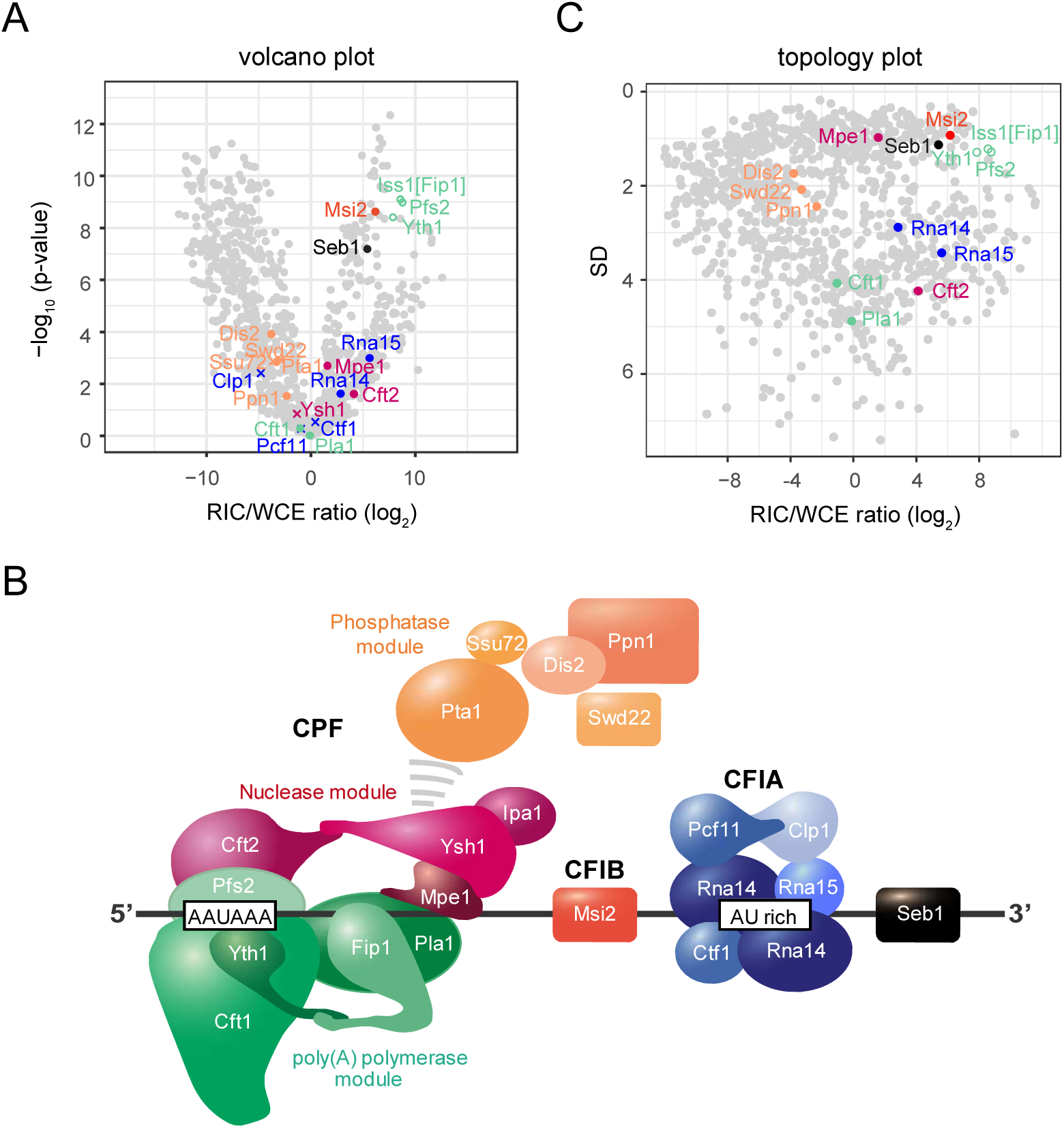
Normalised RIC data captures aspects of multi-protein complex topology. **(A)** Volcano plot of the WCE-normalised poly(A)+ RNA interactome. Components of CFIA, CFIB, and CPF are highlighted. Colour scheme as in B. Full circles denote proteins that were detected in both oligo-d(T) pull-down and WCE, empty circles proteins that were present in the oligo-d(T) pull-downs but not the WCE, crosses proteins that were detected in the WCE, but never in the oligo-d(T) pull-downs (see also 1C). Background values were imputed for proteins without signal in any given sample. **(B)** Putative model showing the organisation of the *S. pombe* 3’ end processing machinery for RNA polymerase II. **(C)** Topology plot of the WCE-normalised poly(A)+ RNA with components of CFIA, CFIB, and CPF highlighted. Standard deviation (SD) was plotted against the fold change of mean MS intensities (log_2_) of proteins recovered from the oligo-d(T) pull-downs of UV-crosslinked samples (3 J/cm^2^) over the input WCE (RIC/WCE ratios; n=6). To obtain SD values, uncertainty propagation was carried out for RIC/WCE ratios with the propagate package and second-order Taylor expansion in R. Only proteins that were detected in the oligo-d(T) pull-down were included in the graph. Background values were imputed for proteins without WCE signal (empty circles).

Subunits of CFIA that recognize the auxiliary regulatory sequences associated with the PAS, Rna14 and Rna15, were also enriched in the oligo-d(T) pull-down. However, other CFIA subunits, Pcf11 and Clp1, as well as the fission yeast specific factor Ctf1, did not crosslink with poly(A)+ RNA. Recombinantly reconstituted CFIA and polymerase module were both found to interact with RNA (Hill et al., 2019). In our RIC experiment CFIA was less enriched on poly(A)+ RNA than those polymerase subunits that bound RNA (Figure 5A), possibly because CFIA associates with RNA elements downstream of the cleavage site. The residual presence of CFIA on poly(A)+ RNA can be explained by previously published data which demonstrates that CFIA can interact with the polymerase module and stimulate poly(A) tail addition (Casañal et al., 2017). Interestingly, the mRNA 3’end processing machinery-associated factor Seb1 (Lemay et al., 2016; Wittmann et al., 2017) was also highly enriched on poly(A)+ RNA (Figure 5A). Seb1 is recruited to the transcription machinery by direct interaction with the C-terminal domain of Pol II and nascent RNA. It preferentially binds RNA at UGUAA sequence motifs, which are frequently present downstream of the cleavage site (Lemay et al., 2016; Schlackow et al., 2013; Wittmann et al., 2017). The enrichment of Seb1in the poly(A)+ interactome suggests that Seb1 also interacts with RNA independently of the distal UGUAA elements. Indeed, we have demonstrated that Seb1 associates with RNA in regions proximal to the transcription start site (Wittmann et al., 2017). All subunits of the phosphatase module interacted with poly(A)+ RNA at low levels only. Our data support a model where the phosphatase module either is not in direct contact with RN, or quickly dissociates from CPF upon cleavage and polyadenylation.

Interestingly, subunits that belong to the same functional module not only possess similar RIC/WCE ratios but also tend to cluster together on the volcano plot. For example, the polymerase module subunit Fip1 clustered together with Pfs2 and Yth1; as did the CFIA subunits Rna14 and Rna15; and the phosphatase module subunits Pta1, Swd22, Ppn1, Ssu72 and Dis2 (Figure 5A and B). To obtain a better spread of the data, yet retain information on data point variance, we plotted RIC/WCE ratios against their standard deviation (SD) (“topology plot”). Consistent with the volcano plot, clustering of components that are part of the same module was observed (Figure 5C). Thus, different submodules of the complex appeared to not only be defined by their degree of association with poly(A)+ RNA (RIC/WCE ratio, x-axis), but to also be characterised by a submodule-typical noisiness of the data (SD, y-axis). To our understanding, this “conservation of noisiness” is an intrinsic property of the RIC method. It is related to the fact that there are systematic aspects to noise in MS, which are primarily connected to signal strength. Hence, the noise of RIC/WCE ratios is influenced by the stoichiometries of RNA-protein interactions. We infer that clustering in the topology plot indicates that a module associates with RNA preferentially as a fully assembled complex rather than through individual non-correlated binding events. We conclude that our RIC-based analysis can provide valuable insights into the organisation of native RNA-regulatory complexes and allow the generation of functional models that can be experimentally tested in the future.

### RNA exosome regulatory factors are RNA-binding proteins

The RNA exosome complex is central to post-transcriptional regulation of various types of cellular RNAs. On its own, the exosome is an unspecific RNA-degrading enzyme with low intrinsic nuclease activity, and therefore requires co-factors to ensure selective and efficient degradation of a variety of different substrates (Kilchert, 2019). Conserved Ski2-like RNA helicases associate with the top of the exosome barrel and are thought to facilitate RNP disassembly and RNA threading through the exosome channel. Fission yeast has two essential nuclear Ski2-like helicases, Mtr4 and Mtl1 (Mtr4-like 1), that localise to the nucleolus and the nucleoplasm, respectively, and mediate degradation of different substrates (Bühler et al., 2007; Egan et al., 2014; Lee et al., 2013; Zhou et al., 2015).

A number of exosome co-factors have been identified through affinity pull-down of the endogenous complexes associated with Mtl1 from fission yeast and Mtr4 from mammalian cells (Tudek et al., 2018). These studies have revealed a similar composition of the exosome regulatory network in fission yeast and higher eukaryotes (but not in *S. cerevisiae*). Although co-factors are required for proper exosome function *in vivo*, it is not entirely clear how they assist in substrate selection and activation of the exosome. One of the co-factors, the YTH domain protein Mmi1, recognises TNAAAC motifs on RNA and mediates exosome targeting to a subset of transcripts (Harigaya et al., 2006; Kilchert et al., 2015; Shah et al., 2014; Yamashita et al., 2012). However, it is not understood how other co-factors contribute to exosome specificity, what regulatory complexes they form, and whether they bind RNA. We plotted known exosome co-factors (Egan et al., 2014; Lee et al., 2013; Zhou et al., 2015) in order to determine how they interact with poly(A)+ RNA (Figure S5A and B). In agreement with Mmi1 being an RBP, it was enriched on poly(A)+ RNA. Iss10, which was shown to interact with Mmi1 (Yamashita et al., 2013), was also enriched in RIC, suggesting that it binds to RNA and might contribute to the specificity of the target selection. The poly(A)-binding protein Pab2 contains an RRM domain and crosslinked well to poly(A)+ RNA. The RRM domain protein Rmn1 and the zf-CCCH type zinc finger protein Red5 were shown to interact with Pab2 by affinity purification and two-hybrid approach (Lee et al., 2013; Vo et al., 2016; Zhou et al., 2015), and both were enriched in the pull-down. Interestingly, the helicase Mtl1 and the Zn-finger protein Red1 both interacted with poly(A)+ RNA and appeared at similar positions in the plot. This is in agreement with the prediction that these proteins form a Mtl1-Red1-Core complex (MTREC), which was proposed based on affinity purification of native complexes (Lee et al., 2013). The role of the other nuclear Ski2-like helicase, Mtr4, in regulating the exosome complex in fission yeast is less well understood.

Together with its interacting partners, the zf-CCHC protein Air2 and the poly(A) polymerase Cid14, it forms a complex that closely resembles the *S. cerevisiae* TRAMP complex (Bühler et al., 2007; LaCava et al., 2005; Vanácová et al., 2005). Fission yeast Mtr4 was proposed to be important for regulating rRNA processing and levels of some non-coding transcripts produced by Pol II, but to play a less prominent role in mRNA regulation than Mtl1 (Bühler et al., 2007; Wang et al., 2008). Surprisingly, components of the Mtr4-Air2-Cid14 complex were highly enriched in on poly(A)+ RNA, which might indicate that this complex plays a broader role in RNA regulation than is currently believed (Figure S5A). Finally, CBC-Ars2, a nuclear cap-binding complex that has been linked to exosome regulation in humans and *S. pombe*, but has widespread functions in nuclear mRNA metabolism (Andersen et al., 2013; Egan et al., 2014; Giacometti et al., 2017), formed a separate cluster with strong RNA-binding activity (Figure S5A). We conclude that most exosome co-factors are RBPs, in agreement with their role in targeting the exosome complex to RNA substrates.

### Comparative RIC with exosome mutants reveals differences in substrate accessibility

The structure of the exosome complex is well described: The core components form a catalytically inert barrel-like structure through which RNA molecules are threaded in 3’ to 5’ direction before they are degraded by Dis3 exonuclease, which is situated at the bottom of the structure (Makino et al., 2013, 2015; Schuch et al., 2014; Zinder et al., 2016) (Figure 6A).

RNA can also be degraded by Rrp6 exonuclease that is located on the top of the exosome. In addition, it has been proposed that RNA can access Dis3 directly without passing through the channel (“direct access”, *da* conformation) (Bonneau et al., 2009; Delan-Forino et al., 2017; Han and van Hoof, 2016; Liu et al., 2014; Makino et al., 2013, 2015). However, the functional contribution of each model as well the mechanisms that direct routing of substrates have remain unexplored. In agreement with the structural studies of the exosome, RIC analysis revealed that the components of the cap region are enriched on poly(A)+, which supports the proposed function of the cap module in RNA selection and recognition (Figure 6B and S5E, in green /yellow). In contrast, proteins that constitute the PH ring of the core showed limited interactions with poly(A)+ RNA (Figure 6B and S5E, in blue). We assume this to be the case because removal of the poly(A) tail from the 3’ end during degradation prevents capture of the crosslinked complex during oligo-d(T) pull-down. In accordance with this assumption, recovery of crosslinked exosome core subunits from HEK293 cells was reported to be much more efficient when crosslinked RNA-protein complexes were enriched by chemical extraction rather than by poly(A)+ RNA selection (Urdaneta et al., 2019).

Behaviour of a given protein in the RIC experiment reflects the mean RNA association of this protein in an ensemble of states. It was suggested that in wild-type, the preferred state of the nuclear exosome is the actively channel-threading conformation (Bonneau et al., 2009; Delan-Forino et al., 2017; Schneider et al., 2012). To test this hypothesis, we sought to perturb the system by using defined mutants of the exosome complex. The Ski2-like helicase Mtl1 in fission yeast has been proposed to be important for exosome-dependent RNA degradation by facilitating substrate RNA unwinding (Egan et al., 2014; Lee et al., 2013; Weick et al., 2018; Zhou et al., 2015). To assess the role of Mtl1 in substrate channelling, we used an *mtl1-1* mutant that carries six point mutations in the area adjacent to the arch domain (Lee et al., 2013) (Figure S5C). Strikingly, crosslinking of Mtl1 protein to poly(A)+ RNA was lost when these mutations were incorporated (Figure 6C), suggesting that the arch domain might contribute to RNA binding of Mtl1. At the same time, Mtl1 protein abundance was not affected in this mutant (Figure S5D). Our data could indicate that RNA binding is important for the function of this essential helicase since the observed loss of RNA binding in this mutant coincided with the stabilization of most known targets of the nuclear RNA exosome (Lee et al., 2013; Zhou et al., 2015 and our unpublished data). In contrast, RNA association of the other exosome-associated helicase Mtr4 was unchanged. Surprisingly, we observed striking differences in exosome crosslinking to RNA in the *mtl1-1* mutant: a pronounced loss of poly(A)+ RNA association with the exosome cap region compared to wild-type (negative shift coefficient in the comparative volcano plot; Figure 6C). This supports a key role for Mtl1 in facilitating engagement of the exosome complex with RNA. Our data support a model where channel-dependent RNA degradation by the exosome relies on the presence of functional Mtl1 and its ability to bind RNA (Figure 6D). At the same time, the decrease in RNA association of the exosome complex was accompanied by a stabilisation of nuclear RNPs (marked by CBC-Ars2, positive shift coefficient; Figure 6C), suggesting that a significant proportion of nuclear poly(A)+ RNA substrates of the exosome depend on Mtl1 for recruitment to the complex. We also noted that RNA association of all cap-associated proteins was equally affected: The ensemble of cap-associated proteins – the cap “signature” – displayed very similar shift coefficients in the *mtl1-1* comparative RIC. Similarly, subunits of CBC-Ars2 shifted as a cluster when comparing wild-type and mutant. This further supports our notion that the detected RNA-protein interactions occur within fully assembled RNP complexes, rather than between RNA and individual, isolated RBPs.

**Figure 6.**
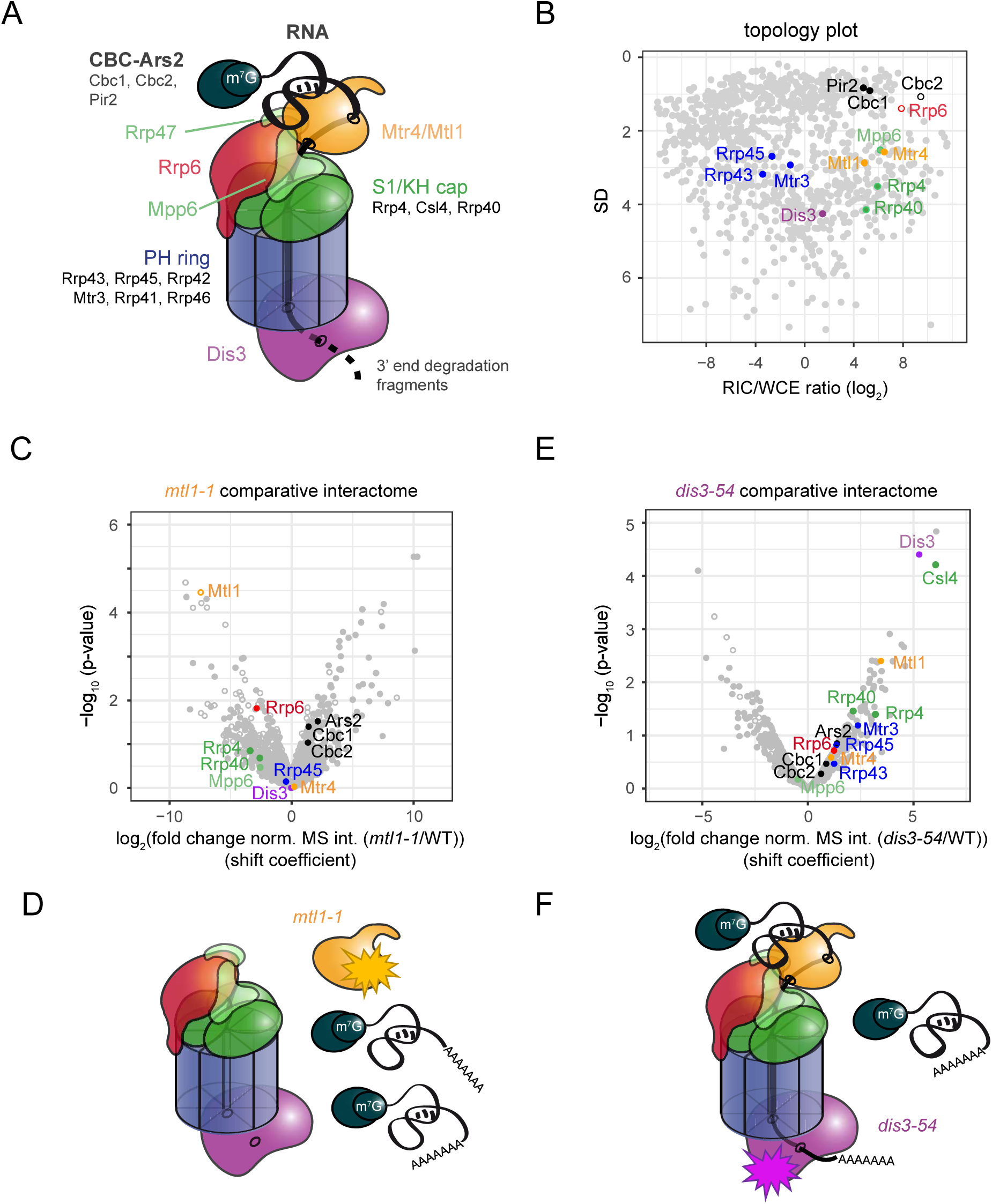
Comparative RIC with exosome mutants captures quantitative differences in RNA channelling. **(A)** Schematics of the nuclear RNA exosome complex, based on crystal structures of the complex (Makino et al., 2013, 2015; Schuch et al., 2014; Zinder et al., 2016). **(B)** Topology plot of the WCE-normalised poly(A)+ RNA interactome with components of nuclear exosome highlighted. Colour scheme as in A. Standard deviation (SD) was plotted against the fold change of mean MS intensities (log_2_) of proteins recovered from the oligo-d(T) pull-downs of UV-crosslinked samples (3 J/cm^2^) over the input WCE (RIC/WCE ratios; n=6). To obtain SD values, uncertainty propagation was carried out for RIC/WCE ratios with the propagate package and second-order Taylor expansion in R. Only proteins that were detected in the oligo-d(T) pull-down were included in the graph. Background values were imputed for proteins without WCE signal (empty circles). **(C)** Volcano plot of a comparative RIC experiment for *mtl1-1*. p-values (-log, moderated Student’s t-test) for the comparison between RIC/WCE ratios of mutant and wild-type interactomes were plotted against the fold-change of RIC/WCE ratios in the mutant vs. wild-type interactome (shift coefficient; n = 3). Components of the nuclear exosome are highlighted. Full circles denote proteins that were detected in the mutant interactome, empty circles proteins that were not detected (but were present in the wild-type interactome). Shift coefficients for individual proteins can be found in Table S3. **(D)** Model: Mtl1 facilitates engagement of the exosome complex with substrate RNA. In the mutant, nuclear RNPs accumulate, but do not engage with the exosome complex. **(E)** Volcano plot of a comparative RIC experiment for *dis3-54* as in C. **(F)** Model: Impairment of Dis3 exonucleolytic activity leads to accumulation of nuclear RNPs, a part of which associate with the exosome complex but fail to be degraded.

We also performed comparative RIC with *dis3-54*, a Dis3 mutant with a Pro to Leu conversion in the RNB domain that has reduced exonucleolytic activity (Murakami et al., 2007). Interestingly, poly(A)+ RNA-binding of the mutant Dis3 protein was strongly increased. We also detected enhanced crosslinking to poly(A)+ RNA all along the exosome barrel and cap (Figure 6E). This could result from the inability of the *dis3* mutant to degrade RNA, leading to continuous association of the channel with polyadenylated RNA, and further supports a prominent role for the channel in RNA decay (Figure 6F).

In addition to its direct function in RNA degradation, Rrp6 was also proposed to play a structural role in anchoring Ski2-like helicases to the exosome (Gerlach et al., 2018; Schuch et al., 2014). Deletion of Rrp6 led to reduced RNA recruitment to the cap region and to Mtl1, which may be explained by the compromised association of Mtl1 with the cap (Figure 7A). We also observed stabilisation of nuclear RNPs in *rrp6Δ*, similar to *mtl1-1.* At the same time, we detected a novel signature in the *rrp6Δ* comparative interactome: a co-shift of Dis3 and Rrp43. Both proteins crosslinked better to poly(A)+ RNA than in the wild-type (Figure 7A) We postulate that this may reflect increased occurrence of the exosome complex in the *da* conformation. In the *da* conformation, Dis3 is rotated to allow RNA substrates to bypass the channel and enter the exonucleolytic centre directly from the surrounding medium (Bonneau et al., 2009; Liu et al., 2014; Makino et al., 2013, 2015). Importantly, substrate RNA that enters Dis3 in the *da* conformation passes Rrp43 on the outside of the PH ring (Figure 7B), which should favour crosslinking of *da* substrates to Rrp43. Hence, the positive shift of the Dis3/Rrp43 signature in the comparative RIC experiment is compatible with an increased use of the *da* route by poly(A)+ RNA in a *rrp6Δ* mutant (Figure 7C). So far, the *da* conformation has only been observed in *S. cerevisiae*, and was suggested to prefer short structured substrates, such as pre-tRNAs and hypomodified tRNA (Delan-Forino et al., 2017; Han and van Hoof, 2016). However, an *S. cerevisiae* mutant that is unable to adopt the *da* conformation – and that has no discernible growth phenotype in a wild-type background – shows a synthetic growth defect with *rrp6Δ* (Han and van Hoof, 2016). Likewise, when we added a C-terminal tag to *S. pombe* Rrp43 (Rrp43-3xHA-3xflag), we observed no growth impairment in a wild-type strain, but synthetic slow growth when the tag was combined with *rrp6Δ* (Figure 7D), possibly because addition of a protein tag at this position interferes with substrate recruitment/access to the *da* route (Figure 7B). Adaptation of the *da* conformation could explain how functional redundancy between Dis3 and Rrp6 is achieved to allow efficient regulation of cellular RNA levels. This would enable Dis3 to execute degradation of Rrp6 substrates in its absence (Delan-Forino et al., 2017).

**Figure 7.**
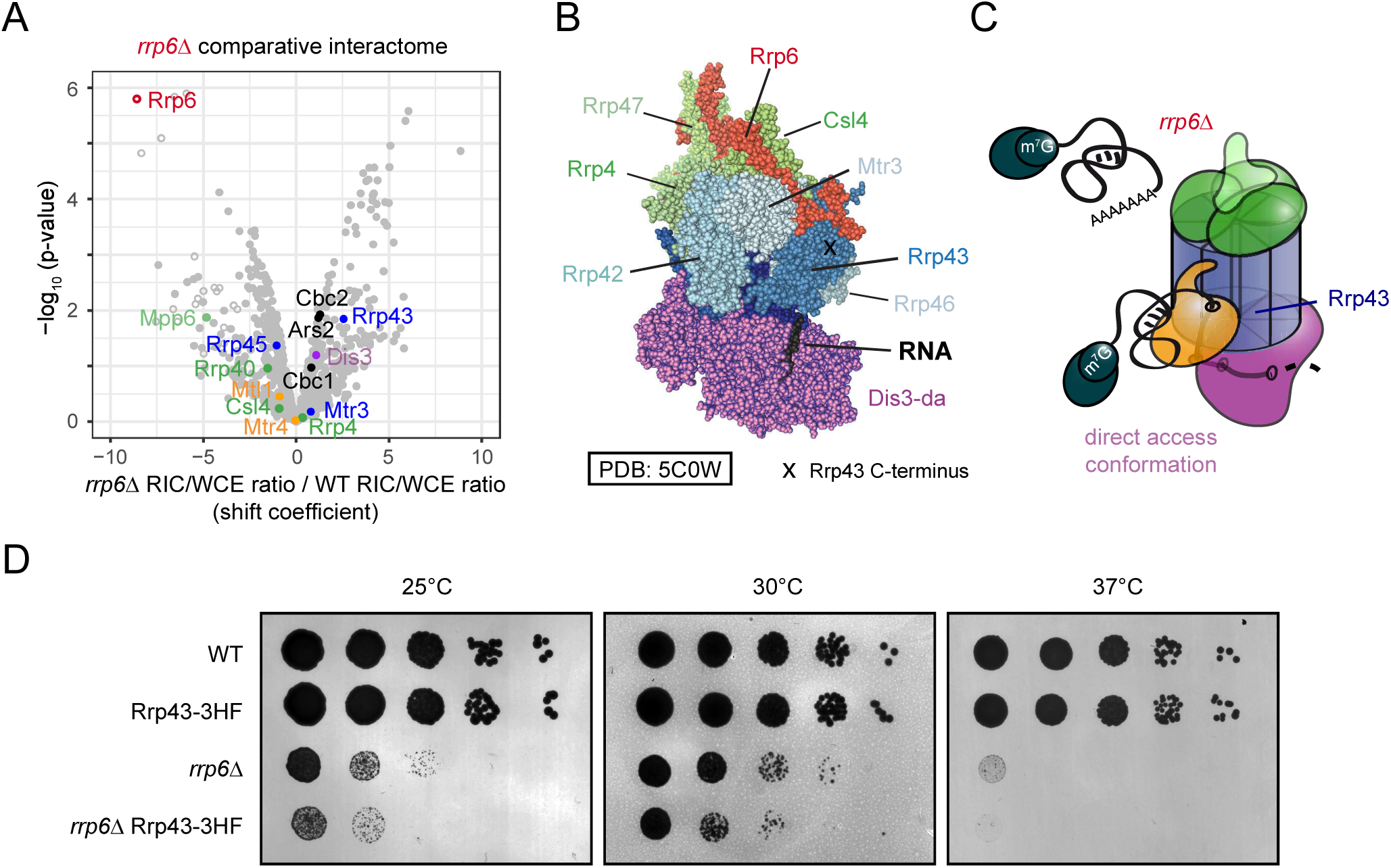
(A) Volcano plot of a comparative RIC experiment for *rrp6Δ*. p-values (-log, moderated Student’s t-test) for the comparison between RIC/WCE ratios of mutant and wild-type interactomes were plotted against the fold-change of RIC/WCE ratios in the mutant vs. wild-type interactome (shift coefficient; n = 3). Components of the nuclear exosome are highlighted. Full circles denote proteins that were detected in the mutant interactome, empty circles proteins that were not detected (but were present in the wild-type interactome). Shift coefficients for individual proteins can be found in Table S3. **(B)** Structure of the *S. cerevisiae* nuclear exosome complex in the direct access conformation (PDB: 5C0W, Makino et al., 2015). Positions of various subunits and of the C-terminus of Rrp43 are marked. **(C)** Model: In the absence of Rrp6, nuclear RNPs accumulate, but fail to engage with the exosome cap region. At the same time, increased RNA crosslinking with Dis3 and Rrp43 indicates a potential rerouting of poly(A)+ RNA substrates from the channelling route to direct access. **(D)** In a plate-based growth assay, addition of a C-terminal tag to Rrp43 results in a slow growth phenotype when combined with *rrp6Δ*, but not in a wild-type background. Serial dilutions (1:10) of the indicated yeast strains were spotted on YES and incubated at the indicated temperatures for several days.

In summary, comparison between wild-type and mutant RNA interactomes allowed us to monitor changes in RNA-protein interactions within the nuclear exosome complex, and to identify preferred routes of substrate access to the complex under different conditions. We propose that comparative RIC experiments will prove very useful to map discrete topological states of large RNP assemblies and allow to assess relative abundances of such states under varying conditions.

### Outlook

To be able to assess relative RNA-binding activities of fission yeast RBPs proteome-wide, we have combined classical poly(A)+ RIC with WCE-normalisation to control for protein abundances. We find that WCE-normalisation offers a distinct advantage: it allows to complement the “snapshot” of the RBPome with robust and quantitative information on the relative enrichment of specific RBPs in the RBPome. Relative enrichment in RIC is influenced by two independent parameters: the degree of association of a given interactor with poly(A)+ RNA, and the UV-crosslinkability of bound RNA and the RNA-interacting region of the protein. For individual RBPs, the relative influence of UV-crosslinkability on enrichment cannot be inferred from the RIC data. However, when we used validated RBPs to benchmark our analysis, enrichment in RIC was almost universal. This has led us to conclude that RIC/WCE ratios can be used as a proxy to estimate *in vivo* RNA-binding activities. We cannot exclude that we may significantly underestimate RNA-binding activities of some RBPs because their mode of interaction with RNA is incompatible with efficient UV-crosslinking.

There are two areas of RNA biology where we consider that having estimates of *in vivo* RNA-binding activities of RBPs can be particularly useful: for the characterisation of (novel) RBPs and RBDs annotated based on proteome-wide screens, and the study of large RNP complexes. RIC and related techniques have proven to be powerful tools in the discovery of RNA-binding proteins (RBPs) and have been employed in different cell types of various organisms to generate comprehensive inventories of their respective RBPomes, and also to identify protein domains capable of interacting with RNA (Baltz et al., 2012; Beckmann et al., 2015; Castello et al., 2012, 2016; Conrad et al., 2016; Kwon et al., 2013; Marondedze et al., 2019; Matia-González et al., 2015; Mitchell et al., 2013; Sysoev et al., 2016). With the identification of novel RNA-binders – and with the recurrent detection in RNA interactomes of proteins with well-described primary functions that are not RNA-related, such as metabolic enzymes – came the debate of how prevalent the observed interactions are in the cell. Here, we have used RNA-binding activity estimates to classify novel RBDs that were identified in previous capture experiments (Castello et al., 2012, 2016). Due to the low number of annotations in *S. pombe*, the assignment of non-classical RBDs to the classes substoichiometric, classical-like, and adaptive is tentative at best, and would greatly benefit from the inclusion of data from other species, since a larger number of annotations would significantly bolster the statistics. Comparison to alternative crosslinking methods, such as formaldehyde crosslinking, would help to remove crosslinking bias. Likewise, the data should be complemented by non-poly(A) RIC data, for which several methods have recently been developed (Asencio et al., 2018; Shchepachev et al., 2019; Trendel et al., 2019; Urdaneta et al., 2019) – this would help to clean up the substoichiometric category of non-poly(A) RNA binders and pave the way for a comprehensive classification of RBDs based on *in vivo* RNA-binding activity. As another caveat, RIC measures enrichment of proteins, not protein domains. For multi-domain RBPs, this can lead to an overestimation of the RNA-binding activity for individual domains. To exclude such multi-domain effects, WCE-normalisation could be combined with the RBDmap approach (Castello et al., 2016).

Although all these limitations apply to the set of substoichiometric RBDs we present here, we consider the attempt worthwhile and a refined classification highly desirable: By revealing whether RNA is the primary bound substrate, or more likely to be a secondary target, it allows to predict the biological role of RBPs.

Recently, very interesting orthogonal approaches have been developed to study RNA-protein interactions, for example R-DeeP, a density gradient centrifugation method that compares sedimentation of protein complexes before and after RNase treatment (Caudron-Herger et al., 2019). R-DeeP allows to quantify the fraction of an RBP that co-sediments with RNA and should therefore be well suited to assess RNA-binding activities for individual RBPs.

Because samples are not crosslinked, we expect R-DeeP to perform particularly well for stable protein-RNA binding events, and maybe less well for RBPs with rapid on-off kinetics. By design, R-DeeP treats multi-protein complexes as one unit and reflects RNA-binding activities of full RNP complexes rather than of individual subunits. In this respect, it differs from RIC, and the information gained from both methods is complementary.

One area where having *in vivo* RNA-binding activities for individual RBPs can be advantageous is the study of multi-protein RNA-binding complex topology. First, it can help to pinpoint subunits that are directly involved in RNA binding within large RNA-protein assemblies, as we have shown for the 3’ end processing machinery. Secondly, it allows to compare the relative RNA-binding activities of different subunits quantitatively. In contrast, CLIP maps RNA-protein interactions at nucleotide resolution but does not allow to assess the efficiency of RNA binding relative to other proteins. Systematic analyses have been carried out on various important cellular complexes involved in gene expression regulation in *S.cerevisiae* (Battaglia et al., 2017). This study has proposed that many proteins associated with transcription form interactions with RNA. In agreement with this study, we report high affinity interaction of the Pol II kinases Cdk9 and Lsk1 with RNA, suggesting that the interaction of Cdk9 and Lsk1 with RNA is not limited to *S. cerevisiae* and conserved in evolution. In mammals, Cdk9 is a component of pTEFb and can be regulated by interaction with specific RNAs (cellular 7SK and HIV-encoded TAR) (Michels and Bensaude, 2008). However, we do not detect poly(A)+ RNA association for most components of PAF complex, nor for the histone modifier Set1, suggesting that their interactions with RNA are likely to be transient and possibly limited to the nascent RNA.

In addition, we demonstrate that even dynamic changes in the RNA binding behaviour of large RNPs can be captured by RIC. Interestingly, we find that different submodules of the same RNA-binding complexes have characteristic signatures in the normalised interactome: They tend to be alike in both relative enrichment and data point noisiness. Due to this, RIC can be very powerful in studying RNP complexes that assemble on RNA differently depending on the condition. If an RNP is dynamic *in vivo*, specific conformations can be defined by a set of conformation-specific RNA-protein interactions. Topology scores derived from a single-condition interactome represent average values for the ensemble of RNP states, and thus do not allow easy identification of conformation-specific RNA-protein interaction signatures. We delimit discrete topological states of dynamic ribonucleoprotein complexes by assessing changes in topology scores caused by specific mutations affecting components of the complex. We demonstrate the power of this approach by monitoring changes in the preferred route of substrate access to the RNA exosome complex in a variety of exosome mutants. We propose that this technique can be employed to identify RNP remodelling events under various conditions and thus provide important insights into the function of the multi-subunit machineries involved in RNA regulation.

## Acknowledgements

We thank S. Grewal and T. Nakamura for strains and constructs. This work was supported by a Wellcome Trust Senior Research fellowship to L.V. (WT106994MA), and a Medical Research Council career development award to A.C. (MR/L019434/1). C.K. is supported by the Emmy Noether Programme of the Deutsche Forschungsgemeinschaft (KI 1657/2-1).

## Methods

### Yeast Strains and Manipulations

All *S. pombe* strains used in this study are listed in Table S4. Standard methods were used for cell growth and genetic manipulations (Moreno et al., 1991). Cells were grown in yeast extract with supplements (YES) at 30°C unless indicated otherwise.

### Poly(A)+ RNA interactome capture (RIC)

Poly(A)+ RNA interactome capture in *S. pombe* was essentially performed as published for *S. cerevisiae* (Beckmann et al., 2015), with the following modifications: *S. pombe* cells were grown in EMMG with limited amounts of uracil (10 mg/l), and labelled with a final concentration of 1 mg/l 4sU for 4.5 h. Cells were grown by the litre, harvested by filtration, then snap-frozen in liquid nitrogen after UV-crosslinking at 3J/cm^2^ in 50 ml PBS. Cells from 3 litres of culture were pooled per experiment and lysed by grinding in liquid nitrogen. RNase inhibitors were omitted from the experiment and all washes after buffer 1 performed at room temperature. The amounts of RNases A and T1 used on the elution fractions before MS were reduced to 1/500 compared to the original protocol. RNase-treated samples were then loaded onto a Vivacon500 filter and denatured in 8 M urea, 100 mM TEAB for 30 min. TCEP was added to a final concentration of 10 mM and the sample reduced for 30 min, then alkylated with 50 mM CAA (final concentration) for 30 min in the dark. The filters were washed with 6 M urea, 50 mM TEAB until the complete removal of detergent. Proteins were digested on filter with 1 µg LysC in 6 M urea, 50 mM TEAB for 4h at 37°C, then with 1 µg trypsin overnight. The flow-through was collected, the filters washed once with 200 µl 0.1% TFA, then with 200 µl 50% ACN, 0.1% TFA, all flow-throughs pooled and dried in a centrifugal evaporator. Samples were resuspended in 50 µL 5% DMSO, 5% FA and analysed on an Ultimate 3000 nanoUHPLC system (Thermo Fisher Scientific) coupled to a QExactive mass spectrometer (Thermo Fisher Scientific). The LC was configured to contain a C18 PepMap100 pre-column (300 μm i.d. x 5 mm, 100 Å, Thermo Fisher Scientific) and an in-house packed analytical column (50 cm x 75 μm i.d. packed with ReproSil-Pur 120 C18-AQ, 1.9 μm, 120 Å) using a 2 h linear gradient (7% to 28% solvent B (0.1% FA in ACN), flow rate: 200 nL/minute). The raw data was acquired on the mass spectrometer in a data-dependent mode (DDA). Full scan MS spectra were acquired in the Orbitrap (scan range 350-1500 m/z, resolution 70,000, AGC target 3xe6, maximum injection time 100 ms). After the MS scans, the 20 most intense peaks were selected for HCD fragmentation at 30% of normalised collision energy. HCD spectra were also acquired in the Orbitrap (resolution 17,500, AGC target 5xe4, maximum injection time 120 ms) with first fixed mass at 180 m/z. A commented version of the protocol is also available (Kilchert et al., 2019). We performed two sets of triplicate experiments of poly(A)+ RNA interactome capture (WT1 + *mtl1-1*; WT2 + *rrp6Δ* + *dis3-54*). The wild-type data sets were merged for analysis of the wild-type interactome, while comparative interactomes were analysed relative to the corresponding triplicate wild-type. A non-crosslinked (noCL) control was included in the first experiment (WT1 + *mtl1-1*).

### Statistical data analysis

The acquired QExactive Raw MS data were processed by the freely available MaxQuant software package (Tyanova et al., 2016). Data were searched against the Uniprot *S. pombe* database alongside as a list of common contaminants provided by the software. The search parameters for the Andromeda search engine (within MaxQuant) were: full tryptic specificity, allowing two missed cleavage sites, fixed modification was set to carbamidomethyl (C) and the variable modification to acetylation (protein N-terminus), oxidation (M). Match between runs was applied. All other settings were set to default leading to a Protein false discovery rate (FDR) of 0.01.

For WCE-normalised interactomes, raw MS intensities were normalised to median = zero for each data set before enrichment was calculated. For all protein data, relative differences were tested against zero with a moderated t-test using the limma package (eBayes function) in R, release 3.32.10, to generate p-values (Ritchie et al., 2015). Proteins detected in both the interactome and the WCE are shown as full circles. In some cases, background values were imputed for WCE normalisation, if no MS intensities were available (open circles). If GO term or Pfam ID annotations are shown, proteins that were not detected in the interactome were also included, as long as they were detected in the WCE; for these, background values were imputed for the interactome signal (crosses). Value imputation was carried out at the level of the triplicate experiments, before the data sets were merged. For the comparative interactomes, WCE-normalised enrichment values for the individual mutants were calculated as before. After WCE-normalisation, uncertainty propagation was carried out with the propagate package and second-order Taylor expansion in R. Data with exact statistics from the error propagation was simulated with mvrnorm (n = 3, empirical = TRUE) from the MASS package and relative differences tested against zero with the limma package. Mutant WCE-normalised MS intensities were corrected by a constant factor to minimise global variance for all crosslinked proteins between wild-type and mutant before calculating fold-change values in the comparative interactomes. Gene ontology enrichment analysis was performed with the enrichment analysis tool on the gene ontology consortium server (release 20181018) (Ashburner et al., 2000; Mi et al., 2017) unless stated otherwise. GO term and Pfam annotations in *S. pombe* were retrieved from EnsemblFungi using the biomaRt package in R, release 2.32.1 (Durinck et al., 2009), information on individual proteins accessed via PomBase (Lock et al., 2018). Boxplots were generated with geom_boxplot(). The lower and upper hinges correspond to the first and third quartiles (the 25th and 75th percentiles). The bold line indicates the mean of the data. The lower and upper whiskers extend from the hinge to the smallest and largest value no further than 1.5 * IQR from the hinge, respectively, where IQR is the interquartile range.

### Expression and purification of Cdk9/Pch1

Full-length *S. pombe* Cdk9 was cloned into one pACE–BacI vector and full-length *S.pombe* Pch1 carrying an N-terminal His-tag followed by a TEV cleavage site was cloned into a second pACE-BacI vector. Purified plasmid DNA (0.5 µg) was electroporated into DH10EMBacY cells to generate bacmids (Berger et al., 2004). Bacmids were prepared from positive clones by isopropanol precipitation and transfected into Sf9 cells (ThermoFisher, UK) grown in Insect-XPRESS (Lonza) with FuGENE HD transfection reagent (Promega) to generate V0 virus. V0 virus was harvested 120 hr after transfection. V1 virus was produced by infecting 50 mL of Sf9 cells grown at 27°C, 300 rpm with V0 virus (2E6 cell/mL, 1:100 (v/v) virus:cells). V1 viruses were harvested 72 hr after proliferation arrest and stored at 4°C. For the co-expression of Cdk9 and Pch1, 500 mL of Sf9 cells (2E6/mL) were co-infected with the two viruses (0.5/100 (v/v) each), cultivated for 72 h at 27 °C, and collected by centrifugation. Cells were harvested by centrifugation (238x g, 4°C, 10 min), resuspended in PBS and centrifuged again, snap-frozen in liquid nitrogen, and stored at −80°C. All subsequent steps were performed on ice or at 4°C. Cells were lysed by sonication in WB buffer (25 mM Tris–HCl pH 8.0, 500 mM NaCl, 10 mM imidazole, 10 mM 2-mercaptoethanol) freshly supplemented with EDTA-free protease inhibitor cocktail tablets (Roche) supplemented with 2500 U of SuperNuclease (Sino Biological Inc.). The lysate was clarified by centrifugation at 230,000g for 40 min. The clarified lysate was filtered through a 0.45 μm filter and applied to a Ni-NTA resin (nickel-nitrilotriacetic acid, 2 mL). The resin was washed with 30 mL WB buffer. Proteins were eluted with 250 mM imidazole. Overnight incubation at 4 °C with AcTEV protease (Invitrogen) was used to remove the N-terminal His-tag from Pch1, and the proteins were run through a Superdex 200 (10/300) gel filtration column (GE Healthcare) equilibrated in column buffer CB (20 mM Hepes 7.5, 150 mM NaCl, 0.5 mM MgCl2, 0.5mM Mg(OAc)2, 1 mM 2-mercaptoethanol). Some fractions contained a contaminant nuclease, others did not. The latter were aliquoted, snap-frozen in liquid nitrogen, and stored at −80°C.

### Cdk9/Pch1 kinase activity assays

Purified *S. pombe* Spt4/5 (Kecman et al., 2018) was used as kinase substrate. End-point reactions were carried out in 40 µl transcription buffer (25 mM HEPES pH 7.5, 40 mM NaCl, 5% (v/v) glycerol, 80 mM KCl, 0.2 mM DTT, 0.25 mM 2-Mercaptoethanol, 0.4 mM MnCl2, 0.5 mM MgCl2, 0.5 Mg(OAc)2, 0.51 mM ATP, 10 µM GTP, 1 µM UTP, 5 ng/µl dsDNA template, 0.5 U/µl of RNasin (Promega)) containing 3.4 µg of Spt4/5 incubated either with buffer or 0.3 µg of Cdk9/Pch1 for 15 min at RT. Reactions were boiled in Laemmli buffer and 1 µg separated on a 7.5% acrylamide gel with 7.5 µM phospho-tag and 50 µM MnCl2. For the time-course experiment, recombinant purified Cdk9/Pch1 (40 pmol) was equilibrated in 20 mM Hepes pH 7.5, 30 mM NaCl, 1 mM DTT, 5% (v/v) glycerol, 0.01% (v/v) NP-40 and 1 mM MgCl2 in a total volume of 30 µL for 20 min at RT. In vitro kinase assays were then performed in a 60 μl reaction with 30 pmol Spt4/5 substrate. MnCl2, DTT and [γ-^32^P]ATP were added to a final concentration of 2.5 mM, 1 mM and 1 μCi., respectively, and reactions incubated at RT. 10 µL were taken for each time point (1, 5, 10, 15, 30 min). The reactions were stopped with LDS-loading buffer and heated at 95°C for 5 min before electrophoresis on 4–12% SDS polyacrylamide gel.

### Fluorescence anisotropy assays with Cdk9/Pch1

Fluorescence anisotropy experiments were carried out at room temperature with 8 nM FAM-labelled RNA of the sequence AUUAGUAAAAUAUAUGCAUAAAGACCAGGC (IDT). The RNA was heated for 5 min at 95°C prior to incubation with proteins. Titration of Cdk9/Pch1 was performed by serially diluting protein with CB buffer (20 mM Hepes 7.5, 150 mM NaCl, 0.5 mM MgCl2, 0.5mM Mg(OAc)2, 1 mM 2-mercaptoethanol) from 1240 nM to 0.32 nM and incubating with RNA substrate for 20 min at RT. Excitation of the ligand was performed with linearly polarized light at 485 nm and emission was measured at 520 nm in parallel and perpendicular planes to the emission plane at 25 °C using a FLUOstar-Omega microplate reader (BMG-Labtech). Each data point is an average of four readings from two different experiments. Anisotropy data were fitted by SigmaPlot software using a standard four-parameter logistic equation to identify Kd:

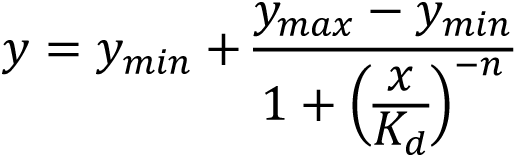

where *y_min_* and *y_max_* are the minimum and maximum anisotropy values, *x* represents the protein concentration, and *n* represents the Hillslope.

---

**Figure S1.**
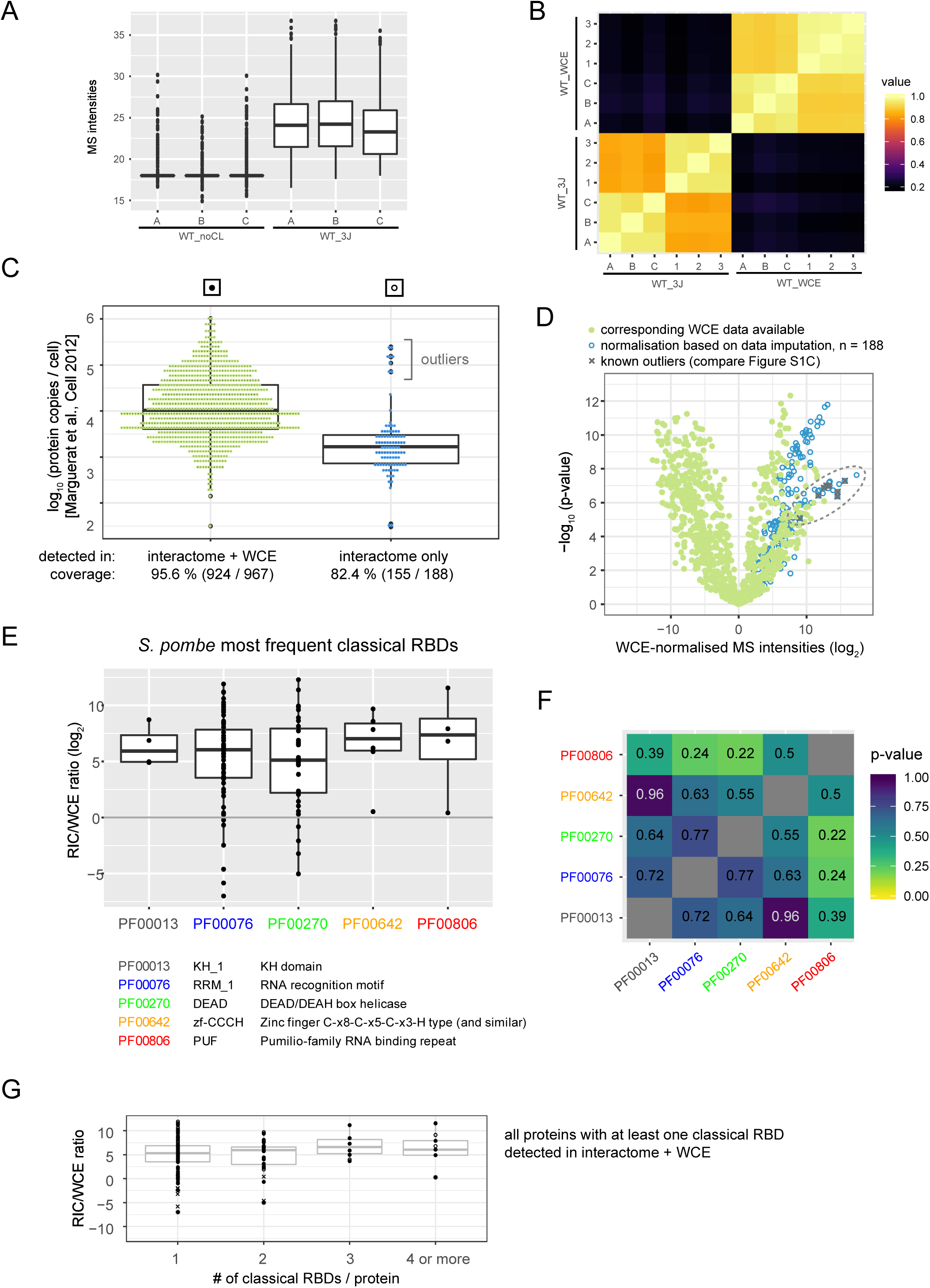
(A) Boxplot of raw MS intensities (log_2_) of proteins recovered from oligo-d(T) pull-downs of UV-crosslinked samples (3 J/cm^2^) versus non-irradiated controls (noCL) (n=3). Few proteins were detected in the noCL control; for all proteins without signal, background values were imputed to be able to calculate the enrichment in the interactome. (B) Spearman’s correlation plot comparing raw MS intensities (log_2_) of proteins recovered from oligo-d(T) pull-downs of UV-crosslinked samples (3 J/cm^2^) and the respective whole cell extracts (WCE) (n = 6). **(C)** Cellular copy numbers for crosslinked proteins that either were (left, green) or were not (right, blue) detected in the corresponding WCE input samples; copy numbers are taken from Marguerat et al. (2012). Coverage designates the fraction of proteins for which copy numbers were available in the Marguerat et al. (2012) data set. On average, cellular copy numbers of proteins that could not be detected in the WCE samples were low compared to proteins that were detected, providing justification for the imputation of background values. Outliers are marked – these are Rpl1201, Hhf2, Eft202, Rps1801, Rps1401, Rps1101, Rps2302, and Rpl1102, and predominantly correspond to ribosomal proteins. 33 crosslinked proteins without WCE signal were not present in the Marguerat et al. (2012) data set and could not be included in the graph. **(D)** Volcano plot, p-values (-log, moderated Student’s t-test) are plotted against the fold change of the mean MS intensities (log_2_) of proteins recovered from the oligo-d(T) pull-downs of UV-crosslinked samples (3 J/cm^2^) over the input WCE (n=6). Crosslinked proteins that either were (green, full circles) or were not (blue, empty circles) detected in the corresponding WCE input samples are indicated. Among proteins without WCE signal, outliers with high cellular copy numbers according to Marguerat et al. (2012) (compare S1C) are designated with crosses. We regard proteins without WCE signal that fall in the region of the plot where these outliers cluster (marked with an ellipse) as low confidence data points. Most low confidence data points correspond to ribosomal proteins (compare also S4C). **(E)** Boxplot of RIC/WCE ratios for proteins with abundant classical RBDs (at least four separate annotations in *S. pombe*). **(F)** p-value matrix for the pair-wise comparison of RIC/WCE ratios for proteins with abundant classical RBDs (at least four separate annotations in *S. pombe*) using a two-sided Student’s t test. **(G)** RIC/WCE ratios for all proteins annotated with at least one classical RBD, separated according to the number of annotated classical RBDs per protein. Full circles denote proteins that were detected in both oligo-d(T) pull-down and WCE, empty circles proteins that were present in the oligo-d(T) pull-downs but not the WCE, crosses proteins that were detected in the WCE, but never in the oligo-d(T) pull-downs (see also 1C). Background values were imputed for proteins without signal in any given sample.

**Figure S2.**
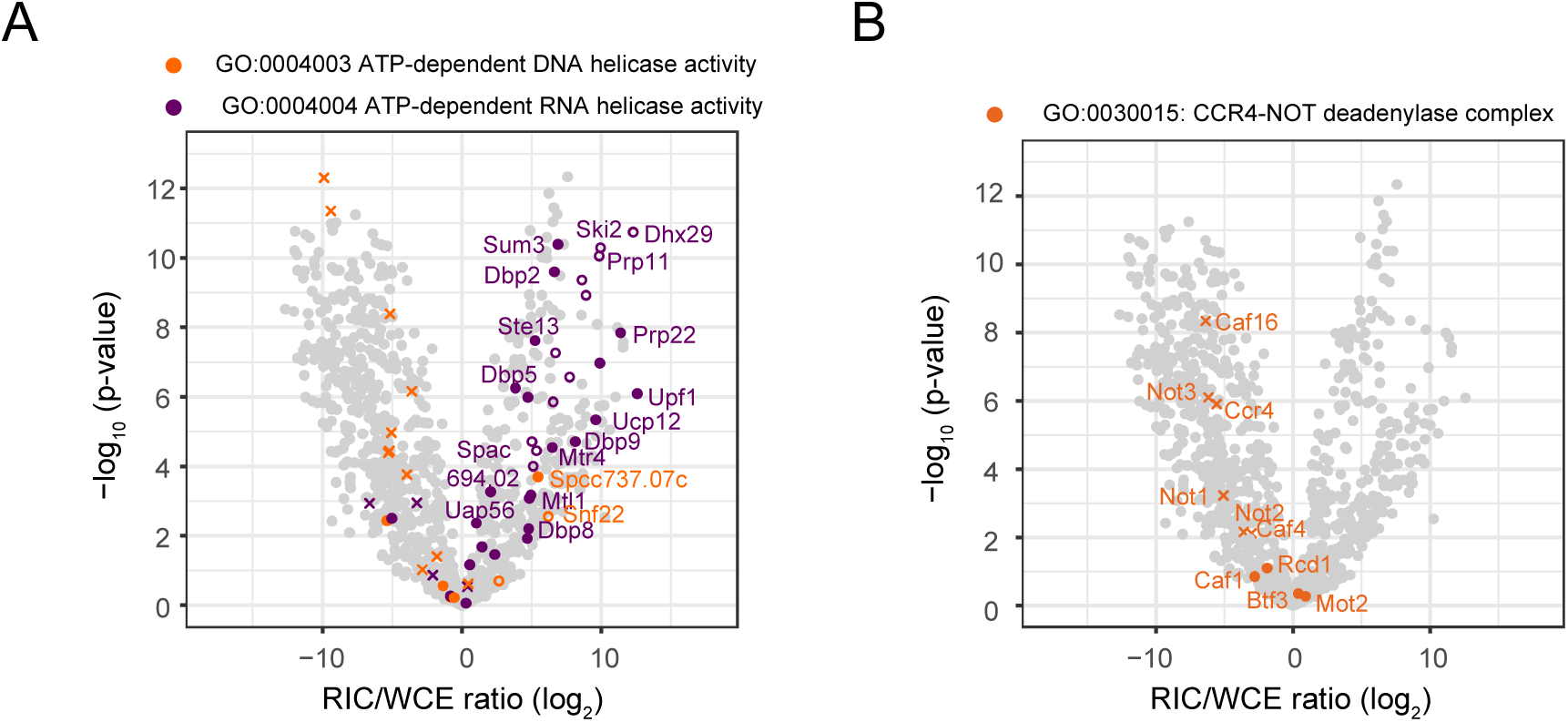
(A, B) Distribution of proteins annotated with various GO terms in the WCE-normalised RNA interactome. Volcano plot, p-values (-log, moderated Student’s t-test) are plotted against the fold change of the mean MS intensities (log_2_) of proteins recovered from the oligo-d(T) pull-downs of UV-crosslinked samples (3 J/cm^2^) over the input WCE (right panel) (n=6). Full circles denote proteins that were detected in both oligo-d(T) pull-down and WCE, empty circles proteins that were present in the oligo-d(T) pull-downs but not the WCE, crosses proteins that were detected in the WCE, but never in the oligo-d(T) pull-downs (see also 1C). In (A), proteins annotated with GO function “ATP-dependent DNA helicase activity” [GO:0004003] or “ATP-dependent RNA helicase activity” [GO:0004004] are indicated in orange and purple, respectively. In (B), proteins annotated with GO component “CCR4-NOT deadenylase complex” [GO:0030015] (orange) are marked.

**Figure S3.**
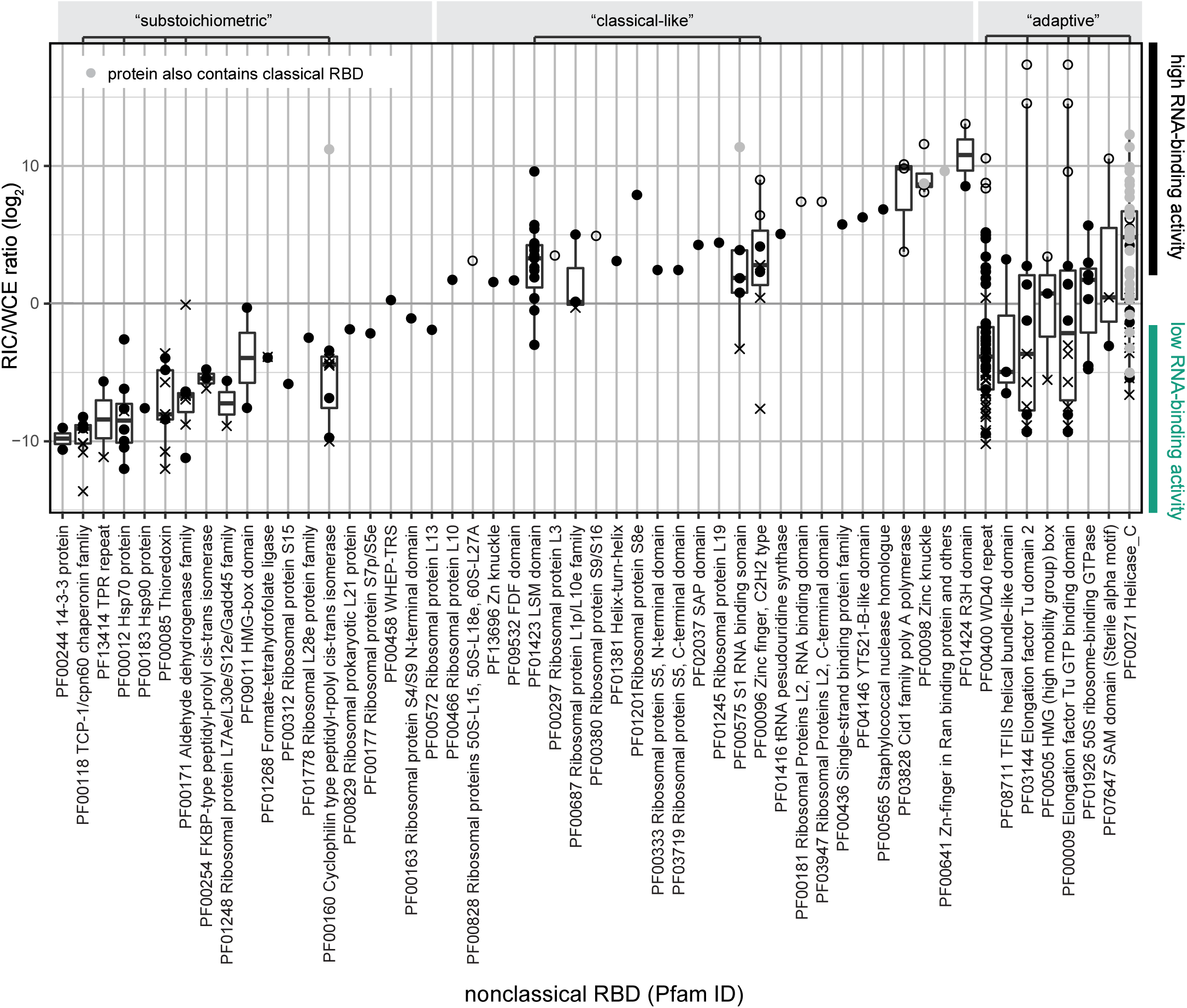
Boxplot of RIC/WCE ratios for all proteins annotated with a non-classical RBD that were detected in the wild-type poly(A)+ RNA interactome capture experiment based on Pfam identifiers given in table S1. Proteins annotated with GO:0022626 [cellular component: cytosolic ribosome] are not included. Full circles denote proteins that were detected in both oligo-d(T) pull-down and WCE, empty circles proteins that were present in the oligo-d(T) pull-downs but not the WCE, crosses proteins that were detected in the WCE, but never in the oligo-d(T) pull-downs (see also 1C). Proteins that also harbour a classical RBD are marked in grey. Abundant non-classical domains (where at least four different domain-containing proteins were detected in the experiment) were classified as either “substoichiometric”, “classical-like”, or “adaptive”, as indicated above the plot.

**Figure S4.**
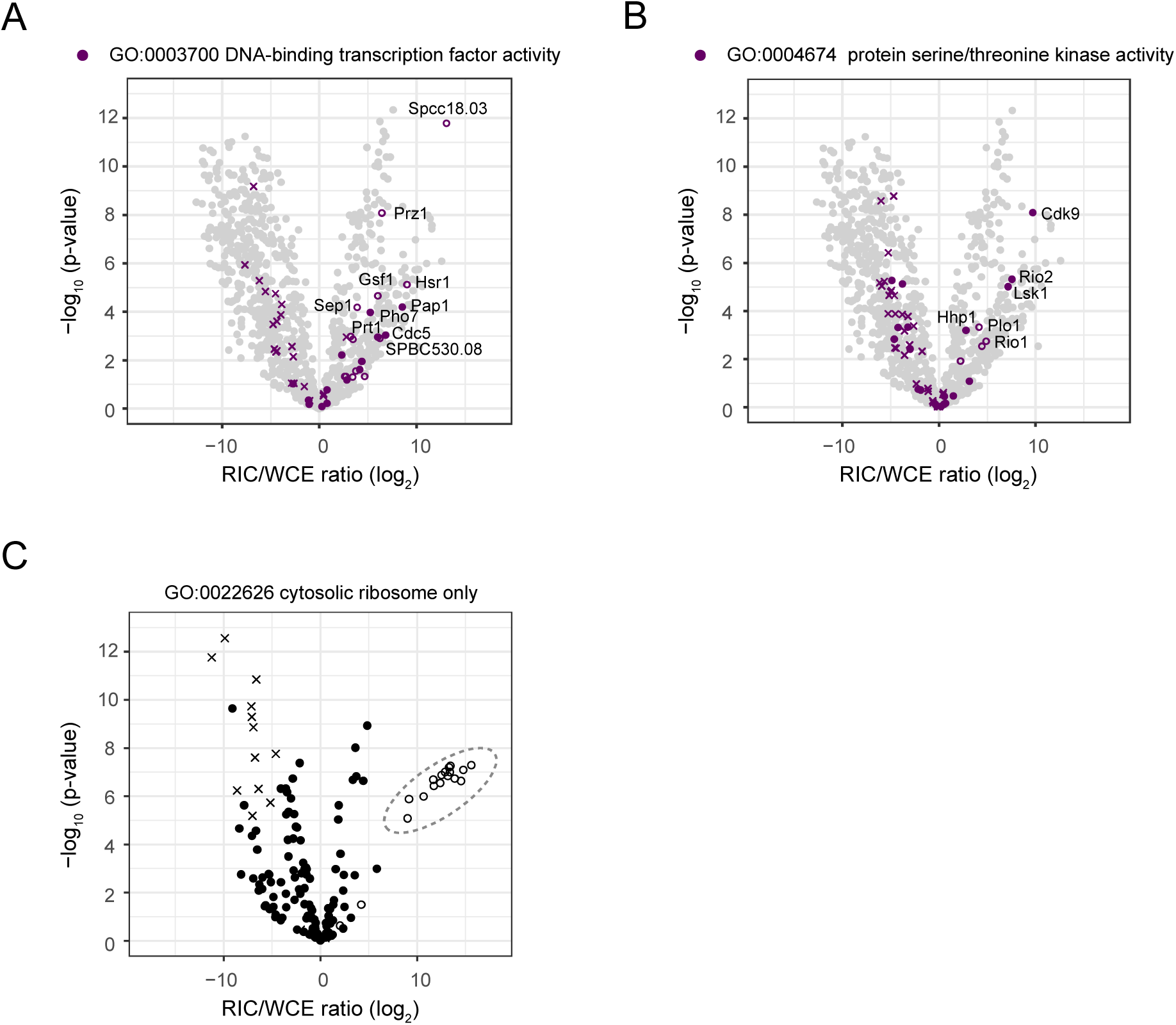
(A, B) Distribution of proteins annotated with various GO terms in the WCE-normalised RNA interactome. Volcano plot, p-values (-log, moderated Student’s t-test) are plotted against the fold change of the mean MS intensities (log_2_) of proteins recovered from the oligo-d(T) pull-downs of UV-crosslinked samples (3 J/cm^2^) over the input WCE (right panel) (n=6). Full circles denote proteins that were detected in both oligo-d(T) pull-down and WCE, empty circles proteins that were present in the oligo-d(T) pull-downs but not the WCE, crosses proteins that were detected in the WCE, but never in the oligo-d(T) pull-downs (see also 1C). **(C)** Distribution of ribosomal proteins (“cytosolic ribosome”, [GO:0022626]) in the WCE-normalised RNA interactome. Volcano plot, p-values (-log, moderated Student’s t-test) are plotted against the fold change of the mean MS intensities (log_2_) of proteins recovered from the oligo-d(T) pull-downs of UV-crosslinked samples (3 J/cm^2^) over the input WCE (n=6). Full circles denote proteins that were detected in both oligo-d(T) pull-down and WCE, empty circles proteins that were present in the oligo-d(T) pull-downs but not the WCE, crosses proteins that were detected in the WCE, but never in the oligo-d(T) pull-downs. RPs with imputed WCE values were among the low-confidence data points (compare Figure S1C and D), and were disregarded in the subsequent analysis.

**Figure S5.**
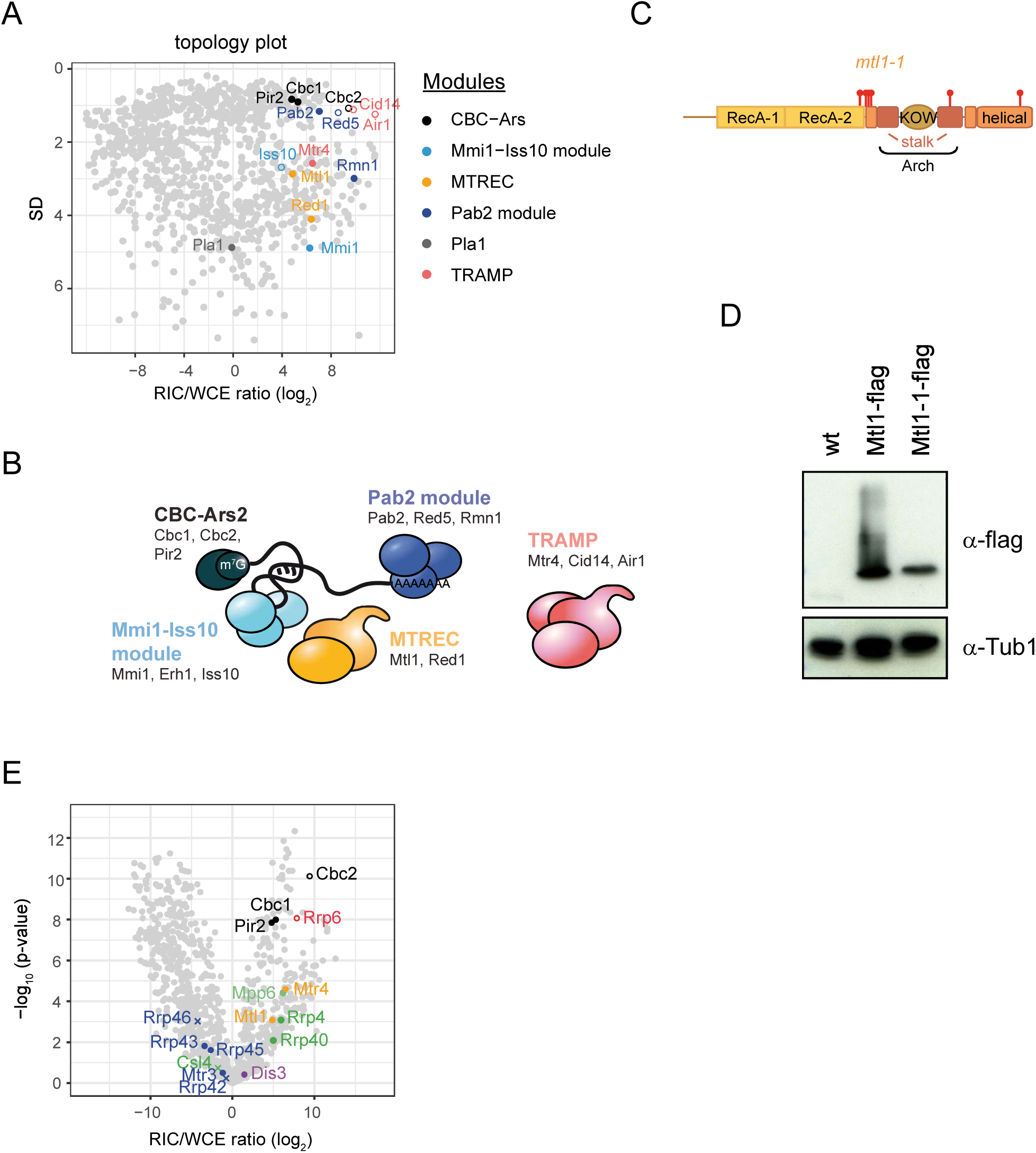
(A) Topology plot of the WCE-normalised poly(A)+ RNA interactome with co-factors of the nuclear exosome highlighted. Colour scheme as in B. Erh1 was not detected in the RIC experiment. **(B)** Co-factors of the nuclear RNA exosome are organised into modules. Schematics based on (Egan et al., 2014; Lee et al., 2013; Shichino et al., 2018; Zhou et al., 2015) **(C)** Domain organisation of Mtl1 protein. Positions of the point mutations the *mtl1-1* mutant are marked as lollipops. **(D)** The mutant Mtl1-1 protein is expressed at similar levels to the wild-type protein; however, the band for the mutant protein is more defined, suggesting that some posttranslational modification may be lost. Equal amounts of yeast lysate were loaded onto SDS-PAGE and analysed by Western blot for flag. An untagged strain was included for reference. Tubulin was used as loading control. **(E)** Volcano plot of the WCE-normalised poly(A)+ RNA interactome. Components of the nuclear exosome are highlighted. Full circles denote proteins that were detected in both oligo-d(T) pull-down and WCE, empty circles proteins that were present in the oligo-d(T) pull-downs but not the WCE, crosses proteins that were detected in the WCE, but never in the oligo-d(T) pull-downs (see also 1C). Background values were imputed for proteins without signal in any given sample.

